# NetrinG1^+^ cancer-associated fibroblasts generate unique extracellular vesicles that support the survival of pancreatic cancer cells under nutritional stress

**DOI:** 10.1101/2021.11.21.469456

**Authors:** Kristopher S. Raghavan, Ralph Francescone, Janusz Franco-Barraza, Jaye C. Gardiner, Débora Barbosa Vendramini-Costa, Tiffany Luong, Narges Pourmandi, Anthony Andren, Alison Kurimchak, Charline Ogier, James S. Duncan, Costas A. Lyssiotis, Lucia R. Languino, Edna Cukierman

## Abstract

It is projected that, in 5 years, pancreatic cancer will become the second deadliest cancer in the United States. A unique aspect of pancreatic ductal adenocarcinoma (PDAC) is its stroma; rich in cancer-associated fibroblasts (CAFs) and a dense CAF-generated extracellular matrix (ECM). This fibrous stroma, known as desmoplasia, causes the collapse of local blood vessels rendering a nutrient-deprived milieu. Hence, PDAC cells are nurtured by local CAF-secreted products, which include, among others, CAF-generated small extracellular vesicles (sEVs). It is well-accepted that upon culturing functionally tumor-promoting CAFs under pathophysiological-relevant conditions (e.g., within self-produced ECM), these cells express NetrinG1 (NetG1) and sustain endosomal pools rich in active α5β1-integrin; traits indicative of poor patient survival. We herein report that NetG1^+^ CAFs generate sEVs that rescue PDAC cells from nutrient-deprived induced apoptosis. Two unique sEVs, NetG1^+^ and α5β1-integrin^+^, were uncovered. The former constitutes cargo of CAF-generated exomeres, and the latter is detected in classic exosomes. Proteomic and metabolomic analyses showed that the sEV-dependent PDAC survival is, at least in part, dictated by the cargo packaged within sEVs in a NetG1-dependent manner. Indeed, despite producing a similar number of vesicles, selected key proteins and metabolites (e.g., glutamine) were incorporated within the unique sEVs. Finally, we found that NetG1 and α5β1-integrin were detected in sEVs collected from plasma of PDAC patients, while their concomitant levels were significantly lower in plasma of sex/age-matched healthy donors. The discovery of these tumor-supporting CAF sEVs opens a new investigative avenue in tumor-stroma interactions and stroma staging detection.

## Introduction

Pancreatic ductal adenocarcinoma (PDAC) is the most common type of pancreatic cancer, accounting for ~90% of cases [1]. PDAC has a sobering 10% overall 5-year survival rate [2, 3] and it is currently the third most deadly cancer in the USA [4]. Since there is a grave need for improved treatment options, it is imperative to better understand the PDAC biology, which encompasses a unique microenvironment with cancer-associated fibroblasts (CAFs) being the most prominent cells. In PDAC, a human tumor mass can on average include 70% stroma, i.e. which primarily includes CAFs and CAF-secreted interstitial extracellular matrix (ECM) [5]. Further, the PDAC tumor microenvironment (TME) undergoes significant restructuring during tumor onset and progression [6, 7], which causes the collapse of local vessels and limits the access to hematogenous sources of nutrients. This stimulates a process of metabolic exchanges that results in the reported TME nourishing of PDAC cells [8–11]. It is therefore important to better understand how pro-tumoral CAFs and their CAF-derived ECMs (CDMs) provide local nutrition support to PDAC cells.

An important mode of paracrine cell-cell communication between CAFs and PDAC cells is through the exchange of extracellular vesicles (EVs). EVs encompass a diverse range of secreted membranous transport-structures that are generated by virtually all cells, are abundant in all bodily fluids, and serve an assorted array of functions [12]. In the context of cancer, EVs are known to regulate various aspects of tumorigenesis, including immune-modulation, metabolic sustenance of cancer cells, and conditioning of the pre-metastatic niche [13–15]. EVs accomplish these functions through intercellular transfer of their cargo, which include a variety of proteins, nucleic acids, and metabolites [15, 16]. Further, EVs contain transmembrane and surface glycoproteins that are integral for facilitating interactions with extracellular molecules [17, 18] as well as plasma membrane (PM)-exposed receptors [19].

This study focuses on small extracellular vesicles (sEVs) in the 20-180 nm range. sEV populations are highly heterogeneous, and include exosomes produced via endosomal routes, and other EVs that are generated via “pinching off” from the PM, known as exomeres and ectosomes [20, 21]. Currently, dissecting the heterogeneous nature of sEV populations constitutes a challenge. This is in part because the unique profile of each sEV population is highly dependent on the type of cell, as well as on the functional state of the cell at the time EVs are generated. As such, the composition of sEVs reflects the functional state of its parent cell, providing sEVs the potential to impart temporal and spatial functions that align with disease stages [22].

We recently identified NetrinG1 (NetG1), a glycosylphosphatidylinositol (GPI)-anchored cell surface protein most commonly observed in the central nervous system for its role in stabilizing glutamatergic synapses [23, 24], as a key driver of pro-tumor CAF functions in PDAC [25]. NetG1 ablation in CAFs modulates the cell’s metabolic secretory profile, and limits the nutritional support provided to PDAC cells [25]. In addition, we previously reported that the PM and intracellular (e.g., endosomal) localization of the main fibronectin ECM receptor, α_5_β_1_-integrin (Int.α_5_), phenotypically and functionally distinguishes tumor-suppressive from tumor-promoting PDAC CAFs, respectively [26]. Importantly, NetG1 expression and Int.α_5_ endosomal enrichment are solely attained *in vitro* if human PDAC CAFs are cultured under pathophysiologic mimicry conditions achieved throughout CDM production [25, 26].

In this study we query the role of NetG1^+^ CAFs in generating pro-tumoral sEVs. We identify a NetG1-dependent sEV function needed to prevent PDAC cell apoptosis induced by nutritional stress. Additionally, we report that normal-like fibroblasts (NLFs), which sustain high levels of Int.α_5_ at the PM [26], fail to express NetG1 and are functionally, albeit not always phenotypically, indistinguishable from NetG1-deficient CAFs. Moreover, we found that tumor supportive CAFs sustained in CDMs secrete sEVs that are packed with significantly higher amounts of metabolites, such as glutamine, than sEVs collected from NLFs. Further, this study describes a novel NetG1^+^ EV particle that is distinct from exosomes. Proteomic and metabolomic analyses uncovered that the sEV-dependent PDAC cell survival advantage is, in part, provided by the type of NetG1-dependent EV cargo as opposed to the amount of vesicles generated. Finally, we provide evidence to suggest that NetG1 and Int.α_5_ are enriched in sEVs harvested from human PDAC patient plasma, yet scarce in plasma obtained from age/sex matched healthy individuals.

Results from this study are significant in that they identified two unique types of tumor-supporting CAF EVs, with evidence of these being detected in patients. Thus, this study facilitates a novel avenue for immersing into the subtleties of the tumor-stroma interactions responsible for PDAC homeostasis and progression, as well as the possibility of establishing future means to detect and monitor dynamic stroma-staging.

## Results

### CDM-dependent pro-tumor human CAFs generate sEVs with unique protein cargo

We previously reported that tumor-promoting human CAFs, cultured within CDMs, sustain PDAC cell survival under nutrient-deprivation in a paracrine manner [25]. We hence tested whether sEVs produced by CDM-dependent pro-tumoral CAFs can account for the reported PDAC survival benefits. We selected to focus on sEVs as this subset includes the most common type of EVs, exosomes, as well as some other vesicles [27, 28]. We employed well-characterized differential centrifugation methods to isolate sEVs, and proceeded to validate the vesicle fractions, using standards dictated by the International Society for Extracellular Vesicles [29]. Using transmitted electron microscopy, we observed that tumor-promoting CAFs [25] generate heterogeneous sized sEVs (Figure 1A). Immunoblots conducted using CDM-sustained CAF lysates and assorted sEV-isolation fractions, indicate that the recovered sEVs include classic/canonical exosome markers like TSG101, CD81, and CD63, and lacked non-specific cellular fragment contaminants such as Histone 3 (Figure 1B). Of note, nanoparticle tracking showed that sEVs sizes were in the reported range, peaking at about 120-180 nm [29] (Figure 1C). Technically, since serum is needed for CDM production and bovine sEVs can be isolated from the media used to generate CDM (e.g., complemented with fetal bovine serum; Figure 1D), CAFs were routinely switched to serum-free media following CDM production, thus assuring that conditioned media (CM) collection included solely CAF-produced sEVs. Following Tandem Mass Spectrometry (e.g., MS/MS) analysis of the CAF sEV fraction, a total of 226 proteins were identified (Supplemental Table 1). Using a cutoff of a minimum of 5 unique peptide sequences per protein, the top 60 proteins were used for a gene ontology query analysis, via Metascape [30]. The most common identified cellular pathways were matrisome-relevant such as, integrin signaling, cell-substrate adhesion, insulin regulation, and wounding (Figure 1E and Supplemental Table 1). Together, these data serve as technical evidence for the isolation of high quality CDM-dependent CAF-generated sEVs, which encompass discrete protein cargo.

**Figure 1:**
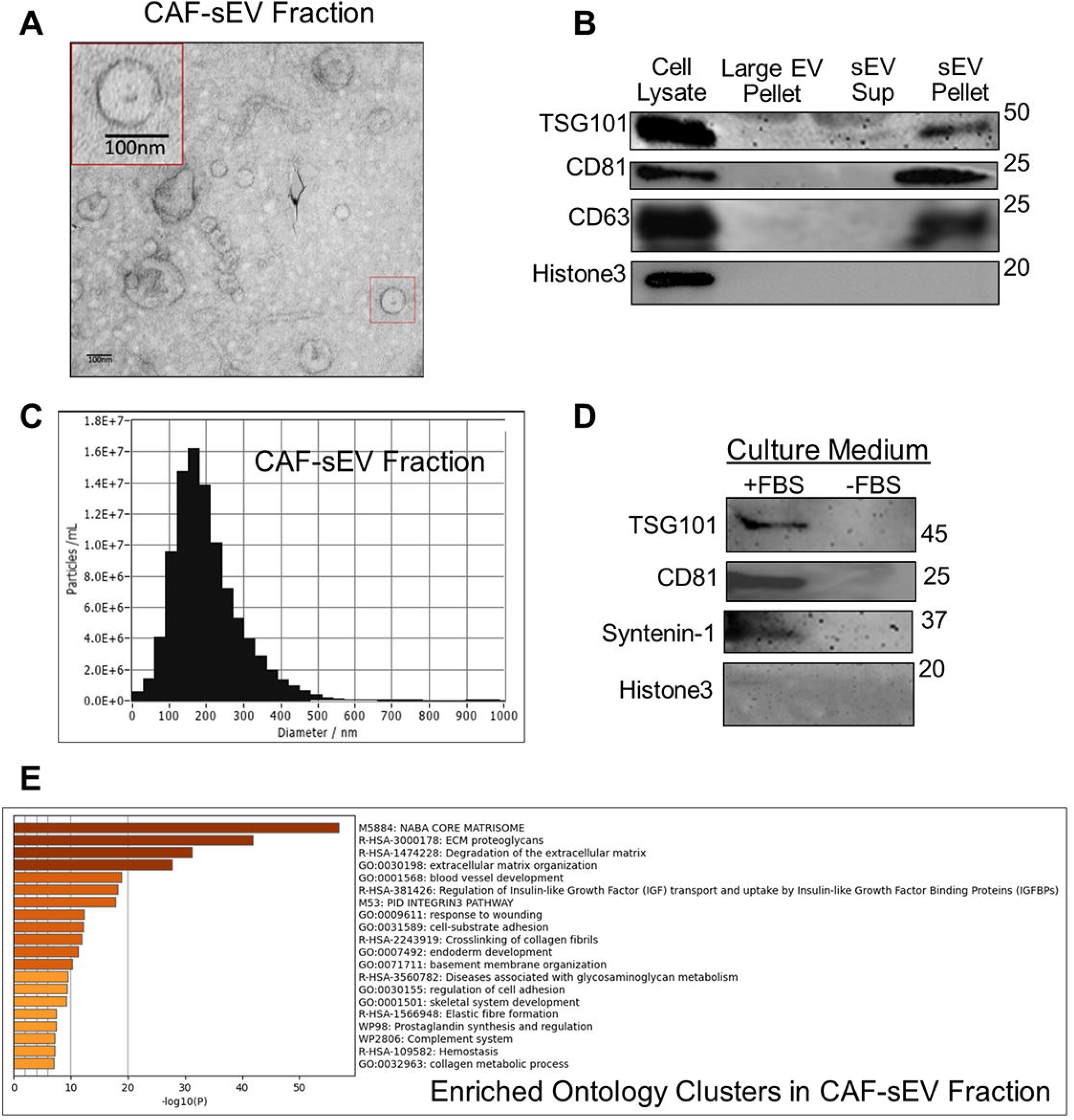
CDM-dependent pro tumoral human pancreatic CAFs generate sEVs with unique cargo. (A) Representative transmission electron micrograph of sEVs isolated from CM collected from pro-tumor CDM-producing human pancreatic CAFs. Red box highlights a zoomed vesicle with canonical exosome-like morphology and size. Scale bar = 100 nm. (B) Fractioned proteins were resolved via SDS-PAGE and immunoblotted. Lysate corresponding to 5 μg protein obtained from the EV-generating CAFs served as loading control. Shown fractions: large EVs pellet (from the 10,000 x *g* centrifuge step); sEV supernatant; and sEV pellet (collected from the 120,000 x *g* centrifuge step (see methods for details). Note that sEV pellet is enriched with canonical exosome markers. (C) Representative Nanoparticle Tracking analysis histogram of sEV pellet fraction. (D) Control Western Blots showing sEVs isolated from media containing FBS (+FBS) and undetected in serum-depleted (-FBS) media. (E) Enriched gene ontology clusters from proteomic analysis of CDM-producing human CAF isolated sEVs, n=3. Note that a full list of CAF sEV proteomic signatures and Metascape pathway analysis can be found in Supplemental Table 1.

### NetG1^+^ CAFs generate sEVs that rescue PDAC cells from nutrient deprivation-induced apoptosis

Well-characterized pancreatic human tumor-suppressive NLFs and tumor-promoting CAFs were used throughout this study [25, 26, 31]. As quality control, and as previously reported, we ascertained that NetG1^+^ pro-PDAC CAFs produce their characteristic dense and anisotropic (e.g., aligned) CDMs, compared to the isotropic (e.g., disorganized) ECMs produced by NLFs (Figure 2A and Supplemental Figure 1). We also validated that, NLFs do not express the canonical contractile fibroblastic marker alpha-smooth muscle actin (αSMA) [32] (Figure 2B), known to be artificially induced in classically-cultured “normal” fibroblastic cells. Note that these types of phenotypic analyses were conducted to assure that sEVs used in all experiments were collected from fibroblastic cells that sustained the known phenotype when cultured *in vitro* [31, 33]. Upon initial analysis of sEVs collected from CAFs vs. NLFs, we noted no significant differences in amounts of sEVs produced by these cells, while minor changes in sEV size distributions were apparent (Figure 2C-D). Importantly, Western Blot analysis of the assorted sEVs revealed an enrichment in the two known pro-tumoral CAF-associated proteins, NetG1 and Int.α_5_ [25, 26], in sEVs isolated from CAFs compared to sEVs from NLFs (Figure 2E). Together, these data demonstrate that while similar amounts of sEVs are generated by the two types of fibroblastic cells, CAF sEVs uniquely include NetG1 and Int.α_5_.

**Figure 2:**
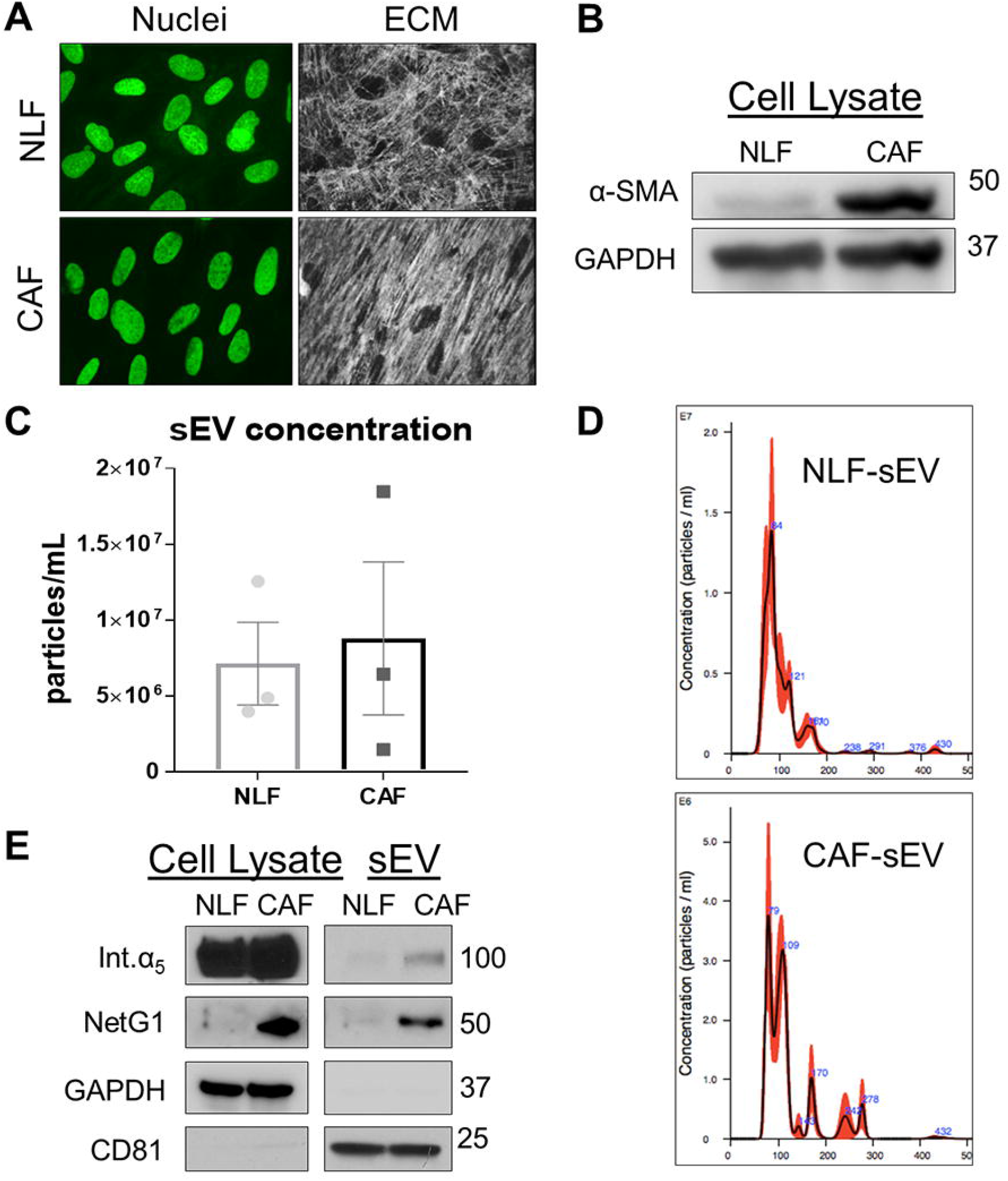
Traits in CAF-generated sEVs differ from the ones in NLFs. (A) Representative confocal immunofluorescent images obtained from NLFs and CAFs. Shown are nuclei (SYBR green) and corresponding fibronectin ECM fibers (white); see Supplemental Figure 1 for measurements. (B) Representative Western Blot denoting αSMA expression in fibroblastic cell lysates. GAPDH = loading control. (C) Aggregated secreted particle concentrations obtained from NLF and CAF sEV fractions showing 3 biological replicates each, using the NanoSight platform. Bars = standard error; p value = 0.9999 using two-tailed Mann-Whitney T test. Note that a comprehensive list of statistical readouts is provided in the Supplemental Table 2 file (Tab = Fig.2C). (D) Representative Nanoparticle Tracking analysis histograms of sEV fractions from NLF and CAFs, using the NanoSight platform. (E) Representative Western Blots denoting NetG1 and Int.α_5_ in ECM sustained cultures of NLF and CAF, cell lysates, and sEV fractions isolated from these. GAPDH = loading control in cell lysate fraction. Note similar CD81 levels in sEV fractions, while NetG1 and Int.α_5_ are enriched in CAF-generated sEVs.

Our previous study demonstrated that NetG1^+^ CAFs provide a survival advantage to PDAC cells when co-cultured within CDMs [25]. Supplemental Figure 2 confirms the ability of the assorted NetG1^+^ CAFs used in this study to sustain tumor-promoting function. Further, we previously showed that NetG1^+^ CAFs can provide this survival benefit in a paracrine manner via CM collected from CAFs [25]. To query whether sEVs isolated from NetG1^+^ CAF CM retain the observed tumor-promoting function, we tested the ability of CAF sEVs to support human PDAC cells cultured under nutrient-deprived conditions (e.g., serum and glutamine free). Similar to CAF-generated CM, for which an obvious survival benefit was observed at 48 hours (Figure 3A), sEVs isolated from CAF-generated CM are about 2-fold more effective than sEVs collected from NLFs in providing a survival benefit to nutrient-deprived PDAC cells (Figures 3B-C, and Supplemental Figure 3A). Of note, prevention of PARP cleavage by CAF sEVs, but not by NLF sEVs, suggest that similar to CM (Supplemental Figure 3B), CAF sEVs prevent apoptotic PDAC cell death induced by nutrient deprivation (Figure 3D). Taken together, these data demonstrate that CAF sEVs protect PDAC cells from nutrient-deprived induced apoptosis.

**Figure 3:**
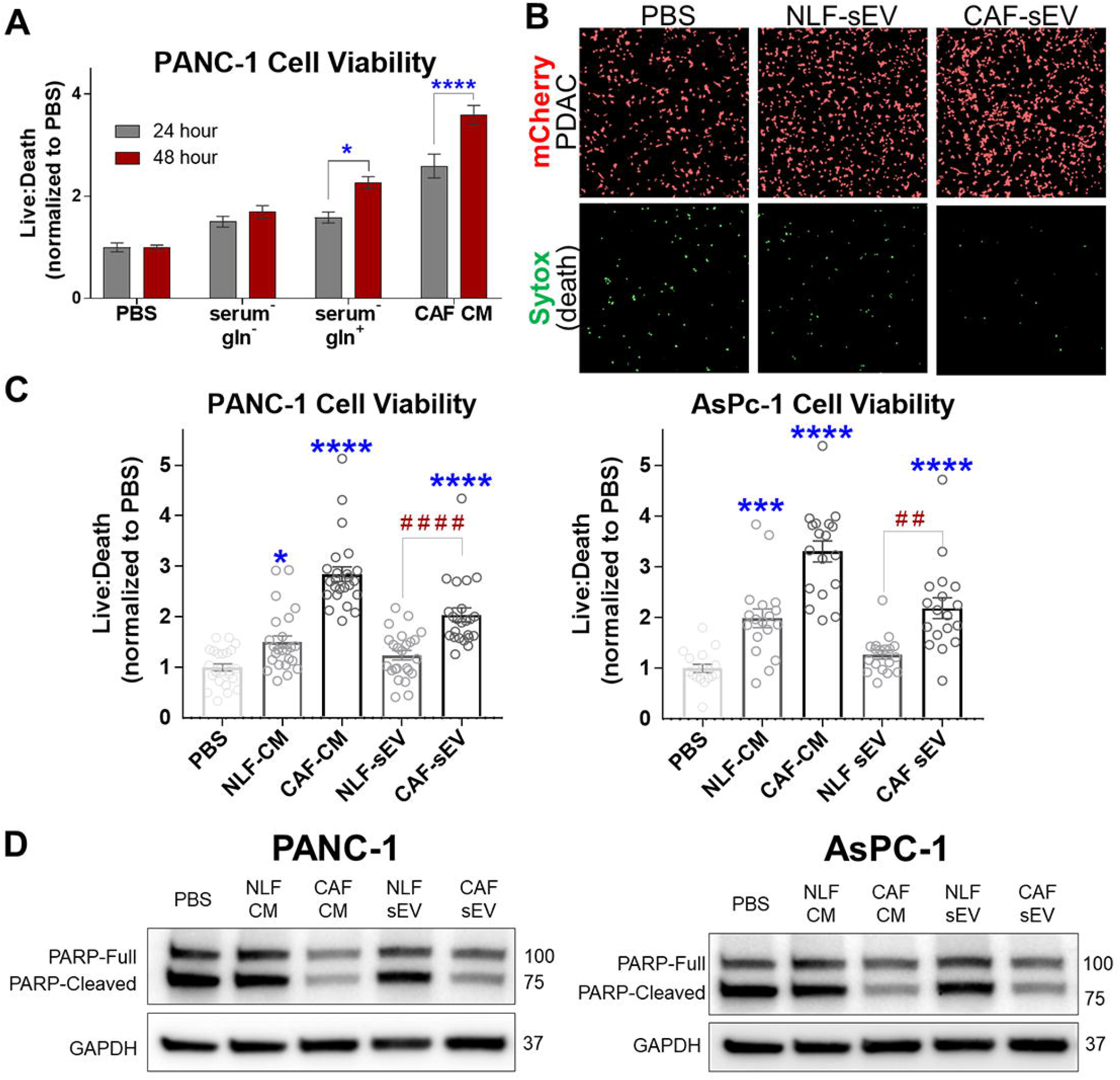
CAF sEVs rescue PDAC cells from starvation-induced apoptosis. (A) Viability Assay comparing 24- and 48-hour endpoints, post-treatment with PBS, starvation media, Glutamine-supplemented starvation media, or CAF-CM. n=3 biological replicates; each replicate consists of 6 technical repeats. All repeats per replicate were normalized to the corresponding PBS-treated condition. Bars = Standard Errors. Statistics: 1-way ANOVA, a minimum of 3 representative fluorescent images were used per condition. (B) Representative Fluorescent images of Panc-1 cells 48 hours post-treatment, used to quantify cell viability. Live cells (shown in red) observed via cellular expression of mCherry, Death (shown in green) observed via Sytox Blue membrane-impermeable nucleic acid stain. (C) Viability assay of PDAC (Panc-1 left; AsPC-1 right) cells at 48 hours gauged as in (A), comparing NLF and CAF CM and sEVs. n=3 biological replicates; each replicate consists of 6 technical repeats. All repeats per replicate were normalized to the corresponding PBS-treated condition. Bars = Standard Error. Statistics: 1-way ANOVA, with multiple comparisons using the Tukey correction. A comprehensive list of statistical readouts is provided in the Supplemental Table 2 file (Tabs = Fig.3A, 3C Panc-1 and 3C AsPC1). * Compared to PBS (negative control), # comparing between conditions noted by connecting lines. (D) Representative Western Blots probing for full-length and cleaved PARP in PDAC cell lysates collected 48 hours post-treatment with PBS, CM, or sEVs from NLF or CAF cells as noted. GAPDH used as a loading control.

### NetG1^+^ CAF-generated sEVs support PDAC cells survival in a NGL1-dependent manner

The sole known heterotypic receptor for NetG1 is the postsynaptic transmembrane protein NetG1-Ligand (NGL1) [34, 35]. We recently reported that NGL1 is expressed in human and murine PDAC cells and that its expression is needed for the nutritional benefit that is imparted by NetG1^+^ CAFs *in vitro* and *in vivo* [25]. Hence, we questioned the levels of NGL1 expression in a panel of human pancreatic tumorigenic cells. Results confirmed that NGL1 levels in these cells are significantly higher than levels detected in benign ductal human neoplastic pancreatic epithelial (HPNE) [36] cells (Figure. 4A). Therefore, we posited a role for the NetG1/NGL1 axis in the observed pro-tumor function from NetG1^+^ EVs. To test this, we generated NGL1 KD Panc-1 cells (Figure 4B) [25], and treated these with NetG1^+^ CAF-generated sEVs as before. Baseline survival during nutrition starved conditions showed that NGL1-deficient cells have limited viability when compared to matched control tumor cells. More importantly, the survival benefit attained when PBS was supplemented with CAF-generated sEVs was abolished in NGL1-KD PDAC cells treated in the same manner (Figure 4C). To further test the involvement of NetG1/NGL1 in this system, we pre-incubated NetG1^+^ CAF sEVs with increasing concentrations of recombinant NGL1 (rNGL1) and questioned if the soluble protein could play an antagonistic role by preventing the observed survival benefit that is imparted by intact sEVs. (Figure 4D) Excitingly, we observed that while rNGL1 produced no noticeable cytotoxicity on its own, if incubated with CAF sEVs, increasing amounts of rNGL1 prompted a significant decrease in PDAC cell survival under nutrient-deprived conditions (Figure 4E). These data suggest that disruption of the NetG1/NGL1 axis reduces the ability for CAF sEVs to nourish nutrient-deprived PDAC cells.

**Figure 4:**
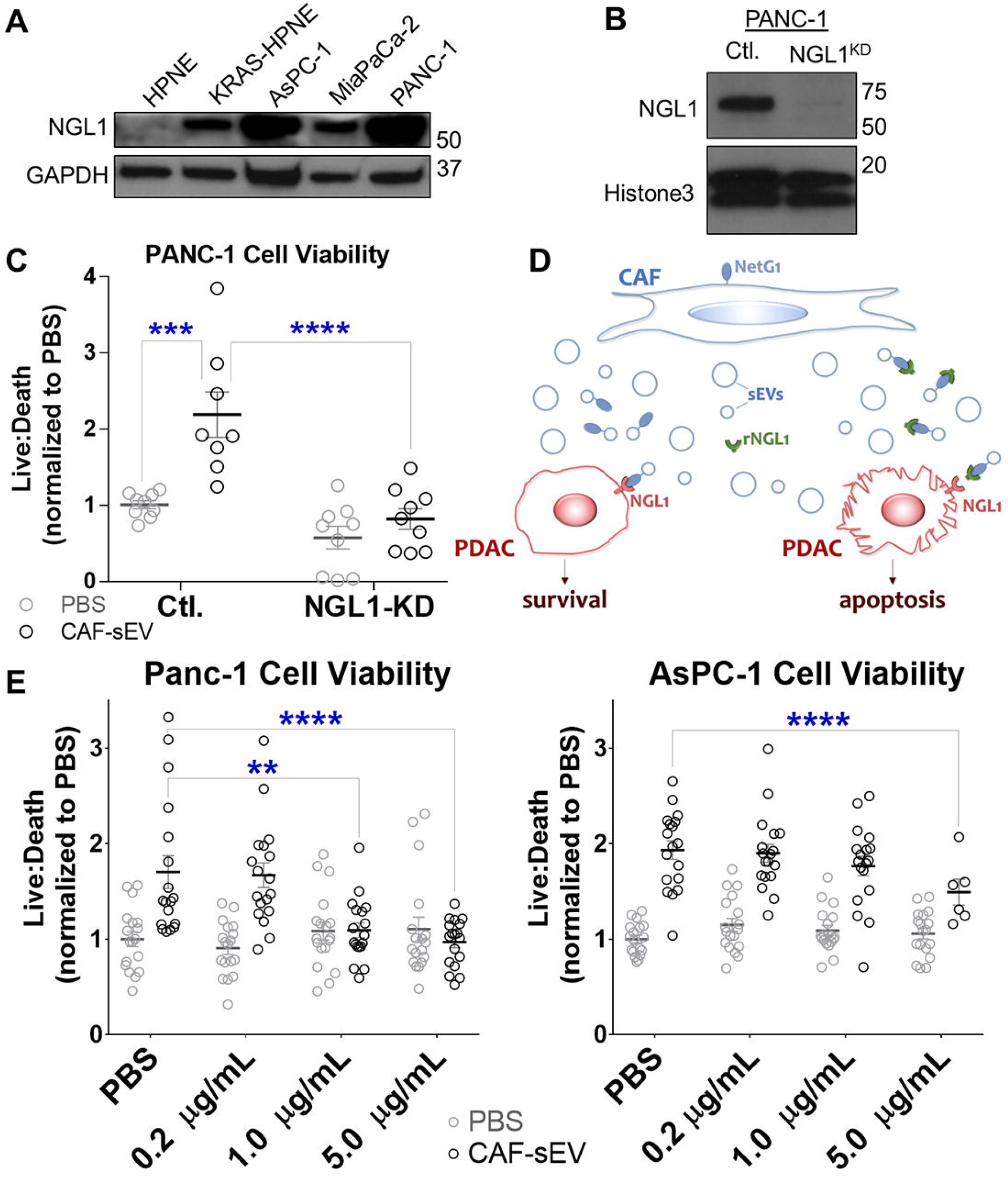
NGL1 expression in PDAC cells is needed for the observed survival benefit imparted by NetG1^+^ CAF-generated sEVs. (A) Representative western blot depicting NGL1 expression in assorted pancreatic epithelial cell lysates. GAPDH was used as loading control. (B) Representative western blot of control vs NGL1-KD PANC-1 cell lysates. Histone H3 served as loading control. (C) PANC-1 viability depicting the ratio of live cells-to-dead cells. (D) Schematic representing the experimental design for 4E; CAF sEVs expressing NetG1 are isolated, incubated with recombinant NGL1 for 20 minutes at RT and then administered to nutrient-deprived PDAC cells expressing NGL1. (E) Dose responsive viability assay of PDAC cells treated with CAF sEVs incubated with recombinant NGL1. For all graphs: n=3 biological replicates; each biological replicate included 3 technical repeats. Technical repeats for each biological replicate were normalized to the PBS-treated condition. Error bars = Standard Error. Statistics = 2-way ANOVA, with multiple comparisons using the Tukey correction. A comprehensive list of statistical readouts is provided in the Supplemental Table 2 file (Tabs = Fig.4C, 4E-PANC1 and 4E-AsPC1)

### NetG1 expression in CAFs is necessary for the EV-mediated survival of PDAC cells

To question the role of NetG1 in the pro-tumor effects noted for CAF sEVs, we used the CRISPRi approach to generate NetG1 knockdown CAFs (NetG1-KD CAFs) [25]. Controls included an empty CRISPRi vector, while CAFs were engineered to overexpress GFP, thus allowing tracking the transfer of “non-specific materials” (Figure 5A). Nanoparticle tracking analysis (NTA) of sEVs showed that just like for NLF cells, NetG1-KD CAFs produce a similar amount of sEVs compared to control CAFs (Supplemental Figure 4A), suggesting NetG1 ablation does not cause a dysfunction in sEV biogenesis. Of note, similar to results obtained when comparing CAF sEVs to NLF sEVs, we saw a change in the average particle size distributions (Figure 5B and Supplemental Figure 4B) suggesting a possible role for NetG1 in modulating the type of EVs that are generated by CAFs. Functionally, and again similar to sEVs derived from NLFs, we observed that sEVs from NetG1-KD CAF lost their ability to protect PDAC cells from starvation-induced apoptosis (Figure 5C-D, Supplemental Figure 4C). Of note, the ability of PDAC cells to uptake sEVs was equivalent, as evidence by the comparable amount of GFP incorporated into PDAC cell lysates following assorted sEV treatment (Supplemental Figure 4D). These results suggested that the type of cargo, as opposed to the amount of material being transferred, is responsible for the observed CAF sEV survival benefit provided to PDAC cells.

**Figure 5:**
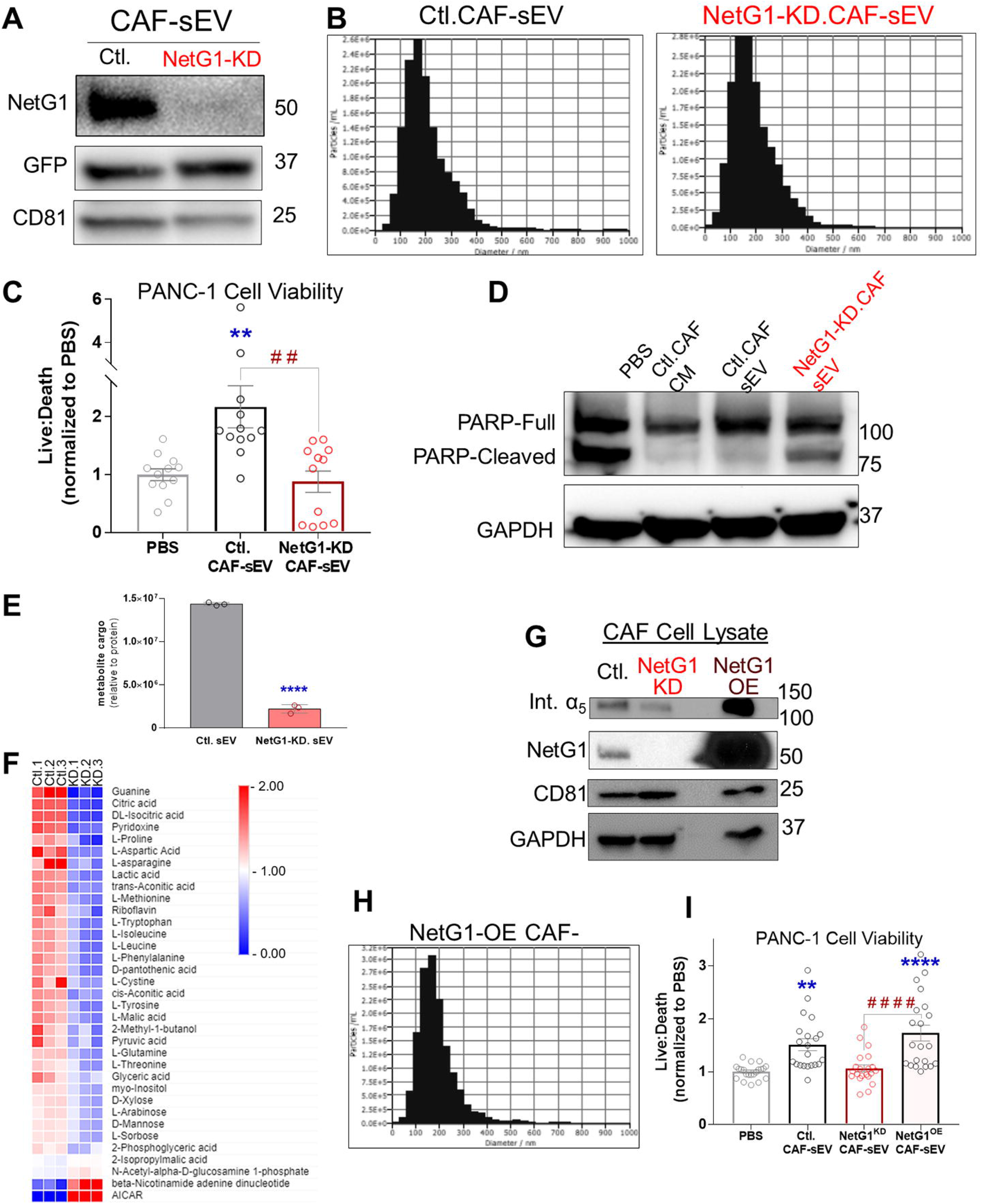
NetG1 expression in CAFs is required for sEV-mediated support of PDAC cells. (A) Representative Western Blots showing effective NetG1-KD in CAF lysates with equal amounts of GFP and CD81. (B) Representative nanoparticle tracking analysis histograms of sEVs from (A), using the ZetaView platform. (C) Viability assay of PANC-1 cells measuring live/dead PDAC cell ratios. n=4 biological replicates; each containing 6 technical repeats. Data were normalized to PBS. Bars = standard error. * compared to PBS (control), and # compared experimental conditions noted by connecting lines. Statistics = 1-way ANOVA, with multiple comparisons using the Tukey correction. (D) Western Blot of full length and cleaved PARP in PANC-1 cell lysates collected 48 hours post-treatment. GAPDH = loading control. (E) Total amount of metabolites detected in assorted sEVs normalized to total protein levels. (F) Heat map depicting control sEV vs KD sEV normalized to median sEV metabolite levels. Shown are the significant products using a t-test p-value of <0.1. (G) Western Blot of cell lysates, validating the levels of expression of the indicated markers in Ctl, NetG1-KD, and NetG1-KD CAFs re-expressing ectopic NetG1 (NetG1-OE) CAFs. GAPDH = loading control. (H) Representative Nanoparticle Tracking analysis histograms of sEV fractions from NetG1-OE CAF, using the ZetaView platform. (I) Viability assay of PANC-1 cells measuring live/dead PDAC cell ratios. n=3 biological replicates; each containing 6 technical repeats. Data were normalized to PBS. Bars = Standard Error. * Compared to PBS (control), # comparing between conditions noted by connecting lines. Statistics = 1-way ANOVA with multiple comparisons using the Tukey correction. A comprehensive list of statistical readouts is provided in the Supplemental Table 2 file (Tabs = Fig.4C, 4I). A full readout of metabolomic analysis can be found in Supplemental Table 3.

NetG1 is known to modulate CAF metabolism, which accounts for the observed ability of CAF-secreted CM to sustain the survival of nutrient-deprived PDAC cells [25]. Hence, to determine if metabolite cargo could explain functional differences in sEVs obtained from control or NetG1-KD CAFs, assorted EVs were boiled and filtered to collect free metabolites, which were then administered to nutrient-deprived PDAC cells. Functional differences between intact vs. boiled fractions were noted, especially with regards to control CAF sEVs (Supplemental Figure 5A). These results justified a follow up assessment of the total amounts of metabolites that are detected in sEVs collected from these cells. Results showed that NetG1-KD CAF sEVs have 6-fold less metabolite content than control CAF-generated sEVs when the measured metabolite content was normalized to sEV protein (Figure 5E). Further, when comparing the types of metabolites included in the assorted sEVs, several specific metabolites were significantly enriched in control CAF sEVs, including glutamine as previously reported [25] (Figure 5F). Pathway enrichment analyses highlighted that upon NetG1 loss, CAFs generate sEVs with cargo that is significantly deficient in numerous key metabolic pathways such as, ammonia recycling, citric acid cycle, transfer of acetyl groups into mitochondria, the Warburg effect, the Urea cycle, and more (Supplemental Figure 5B). These results suggest that loss of NetG1 in CAFs reduces the pool of available nutrients in CAF-generated sEVs, leading to a decrease in PDAC survival under nutritional stress.

To further validate the reciprocal role that NetG1^+^ CAF sEVs play in protecting PDAC cells from nutrition-deprived induced death, we re-expressed NetG1 in NetG1-KD CAFs (NetG1-OE CAF). NetG1 overexpression was confirmed by Western Blot (Figure 5G). Similar to NetG1 KD, overexpression of NetG1 did not change total sEV numbers yet the sEV size distribution was again slightly shifted (compare Figure 5B to 5H). Notably, the re-introduction of NetG1 effectively re-instituted the ability of sEVs to provide a significant survival benefit to PDAC cells cultured under nutrient-deprived conditions. In fact, survival levels observed were similar to those attained from control CAF sEVs (Figure 5I and Supplemental Figure 4E). Taken together, these data demonstrate that CAFs require NetG1 expression to produce sEVs capable of successfully supporting the viability of PDAC cells under nutrient deprivation.

### Distinct NetG1^+^ nanoparticles differ from Int.α_5_^+^ exosomes and together sustain PDAC cell survival

Knowing that EV biogenesis is dictated by the cellular location of the proteins that are sorted into the EVs at time of EV production, where PM and endosomal proteins are differentially distributed among assorted EVs [37], we next confirmed the suspected PM localization of CDM-dependent NetG1 expression in CAFs, as well as the reported endosome-to-PM relocation of Int.α_5_ in NLFs compared to NetG1^+^ CAFs [26] (Figure 6A). Further, and supported by Figure 1A, it is also well-known that sEV fractions are highly heterogeneous [38]. These facts together with results shown in Figure 2E demonstrating that both NetG1 and Int.α_5_ constitute sEV cargo, prompted the hypothesis that the two key proteins could be sorted into different vesicles. To test this premise, we conducted a series of double immuno-labeling electron microscopy experiments, using gold or quantum-doted conjugated primary antibodies. The tetraspanin CD81 was used as a canonical exosome biomarker and its location was compared to that of NetG1 and Int.α_5_. In line with our hypothesis, NetG1 was detected in distinct nanoparticles (DNPs), also known as “exomeres” [21, 39], spanning about 20 nm in diameter and not on CD81^+^ vesicles, or on EVs of sizes similar to exosomes (e.g.~100 nm). In comparison, Int.α_5_ co-localized with CD81^+^ EVs and failed to be detected on exomere-like DNPs (Figure 6B).

**Figure 6:**
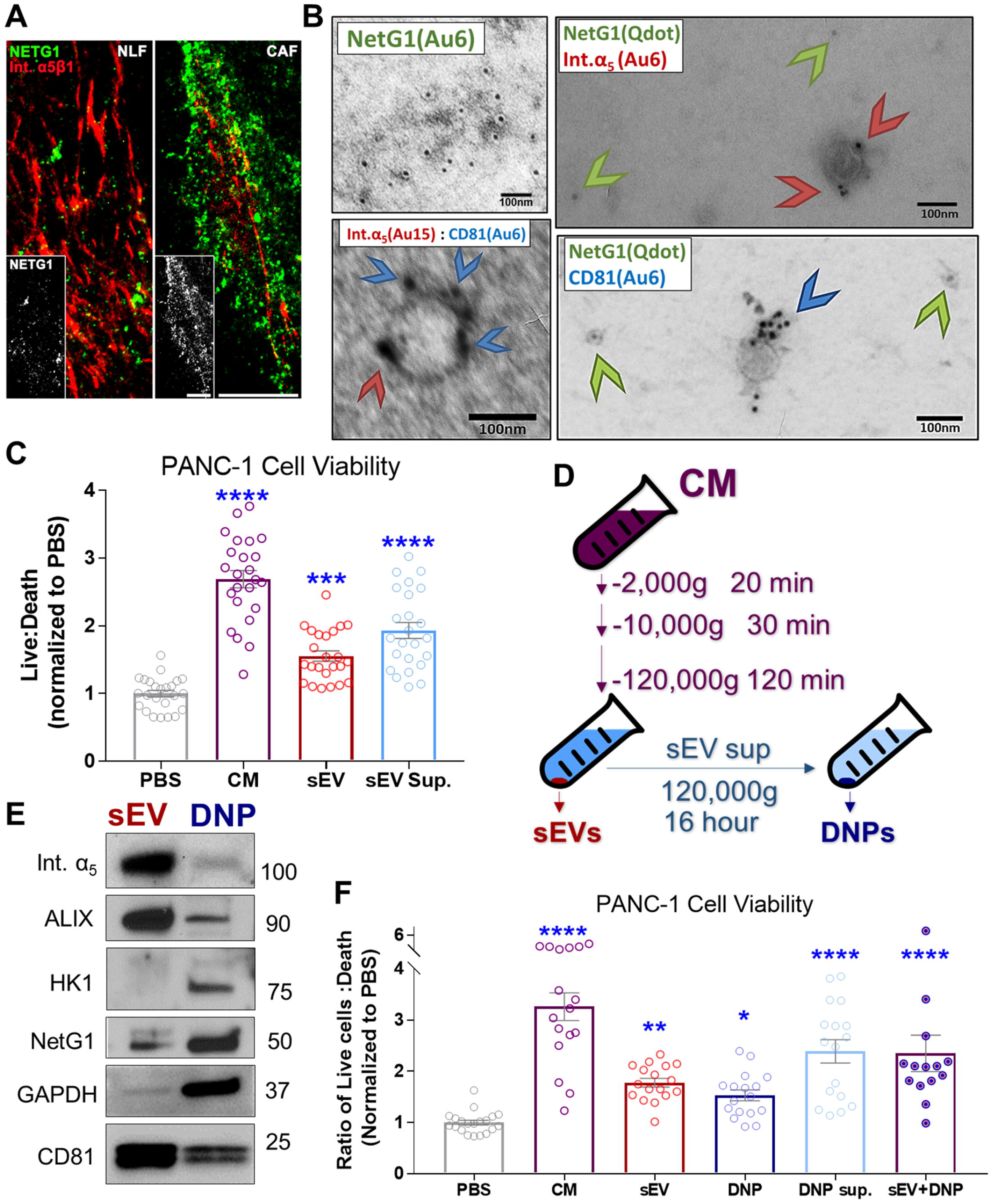
NetG1 is enriched in distinct nanoparticle (DNP) sub-populations which support PDAC viability. (A) Double indirect immunofluorescence tagging NetrinG1 (NetG1; green) and Int.α_5_β_1_ (Int.α_5_; red) on the PM of NFL (left) vs. CAF (right) cultured within self-generated ECMs. Note that the experiment was conducted under non-permeable conditions designed to detect PM localized and extracellularly exposed epitopes preventing antibodies from penetrating to intracellular locations. Scale bars = 10 μm. Inserts are monochromatic images of NetG1. (B) Representative TEM images of immunolabeled CAF sEVs. (Upper left) Immunolabeling of NetG1 with anti-NetG1 antibody-conjugated gold particles (black dots), (upper right) double immuno-detection of NetG1 and CD81 using anti-NetG1 antibody-conjugated quantum dot (low density dots; green arrowheads) and anti-CD81 antibody-conjugated gold particles (high density dots; red arrowheads), (Lower left) double immune detection of CD81 (small dots pointed by blue arrowheads) and activated Int.α5 using SNAKA51-conjugated gold particles (big dots pointed by red arrowheads). (Lower right) double immuno-detection of NetG1 and activated Int.α5 using anti-NetG1 antibody-conjugated quantum dot (low density dots; green arrowheads) and SNAKA51-conjugated gold particles (high density dots; blue arrowheads). (C) Viability assay of PANC-1 cells treated with the indicated fractions. (D) Schematic describing the modified differential ultracentrifugation technique used to obtain the fractions used in subsequent experiments. (E) Representative Western Blot depicting NetG1 and Int.α_5_ levels in sEV and matched DNP fractions. CD81, ALIX = positive sEV controls; HK1, GAPDH = positive DNP controls. (F) Viability assay of PANC-1 cells treated with the indicated fractions. For F and C: n=4 biological replicates; each counting with 6 technical repeats. Repeats per replicate were normalized to PBS condition. Bars = Standard Error. * Compared to PBS. Statistics = 1-way ANOVA, with multiple comparisons using the Tukey correction. A comprehensive list of statistic readouts is provided in the Supplemental Table 2 file (Tabs = Fig.5C, 5F).

Because the sEV supernatant fraction is known to be enriched in DNPs/exomeres [40], we next determined whether this fraction contained any additional pro-tumor functions. Interestingly, the sEV supernatant still contained a significant amount of pro-tumor activity (Figure 6C), suggesting that this fraction could include a unique population of pro-tumor DNPs (e.g., exomeres). To further fractionate subpopulations of EVs from CAF-CM, we employed a modified version of the differential ultracentrifugation protocol [40] reported to effectively precipitate DNPs from sEV-supernatants (Figure 6D). Importantly, we confirmed that the DNP fraction was indeed enriched with smaller particles than the sEV fraction (Supplemental Figure 6). Excitingly, we noted an enrichment in NetG1, in the DNP fraction, which was accompanied by a decrease in Int.α_5_ and other known exosome markers (e.g., ALIX and CD81). Of note, glycolysis proteins (e.g., HK1 and GAPDH) that have been previously reported to be expressed in DNPs [20, 40] were instead enriched (Figure 6E). Of note, the NetG1 enrichment observed in DNPs was also functionally significant in imparting a survival benefit to PDAC cells cultured under nutritional stress as before (Figure 6F). These results are in agreement with out hypothesis that NetG1 and Int.α5 could identify distinct populations of sEVs (ie.g., exomeres vs. exosomes), and that each population could impart a distinctive functional impact on PDAC survival.

### Proteomic and metabolomic profiling of distinct CAF-EV subpopulations suggest unique roles in pro-survival function

In order to query the specific protein differences between the sEV and DNP fractions, we conducted a Label Free Quantitation (LFQ) protein profiling [41]. Principal component analysis of the sample replicates indicated that the fractions corresponded to the enrichment of two distinct populations and presented with a high degree of consistency within experimental conditions (Figures 7A). The distinct protein enrichments per particle type was visualized via a heat map and was accompanied by a volcano plot representation of the data, which identified representative proteins for each of the two fractions (Figures 7B and 7C), while the Venn Diagram (Figure 7D) highlights the enrichment of selected proteins for each of the two types of EV fractions. Consistent with reported data and with our above-presented results, the sEV fraction was significantly enriched in known canonical exosome markers such as CD81, CD63, ALIX, and Syntenin1. This fraction also included β1-integrins (known to form heterodimers with Int.α_5_), as well as HLA-class1 histocompatibility antigen, and beta-actin, as previously reported [42]. Further, identified proteins involved in glycolysis and other relevant metabolic pathways were enriched in the DNP fraction, including the previously reported lactate dehydrogenase A [43], transketolase [44], and beta-hexosaminidase, the last of which has also been reported in unique DNPs known as “exomeres” [21, 45]. Next, using Metascape’s gene annotation and analysis resource [30], we identified a number of signaling pathways unique to each fraction (Supplemental Figure 7A). Metabolomic analyses comparing sEVs and DNPs (normalized to total EV protein), indicated that CAF sEVs incorporate 9-fold more metabolite cargo than CAF-generated DNPs (Figure 7E). In terms of specific metabolites, querying these factors normalized to median metabolite levels, sEVs included commonly known pro-tumor metabolites such as glutamine and proline, while only 3-methylglutaric acid was enriched in DNPs (Figure 7F). Regarding the specific metabolic pathways identified as EV cargo, in comparison to DNPs, sEVs were enriched in mitochondrial substrates, sugars, and amino acids (Supplemental Figure 7B). These results suggest that the observed pro-survival function associated with DNPs is mostly due to non-metabolite cargo while sEVs’ cargo included significant amounts of unique pro-tumoral factors (e.g., metabolites). Together, these data provide evidence of the distinctive properties proposed for sEV and DNP fractions isolated from CAFs and suggest individual functions for the assorted EVs.

**Figure 7:**
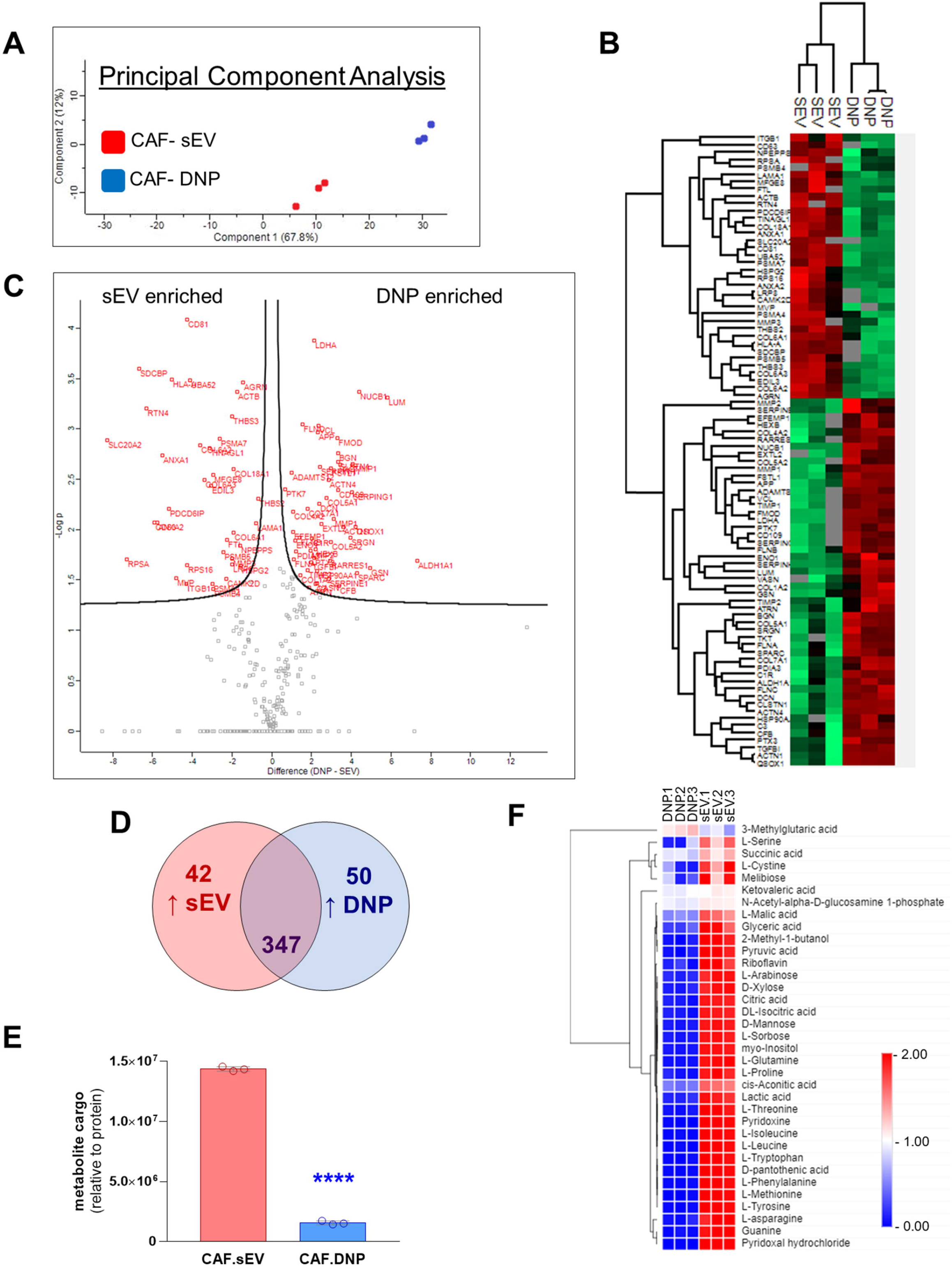
Proteomic and metabolomic profiling of CAF sEVs vs DNPs reveals differences in their cargo. (A) Principal component analysis of sEV and DNP replicate samples. Tight clustering of replicates indicating reliable reproducibility in sample preparation and processing. (B) Heat map visualizing protein enrichment in individual replicates of sEV and DNP fractions. (C) Volcano Plot comparing individual protein levels in sEV vs. DNP fractions. Red labels indicate statistically significant proteins enriched in sEV (left side) or in DNP (right side) fractions. (D) Venn diagram visualizing the numbers of unique proteins enriched in sEV or DNP fractions. Parameters for student’s t-test were as follows: S0=2, side both using Benjamini-Hochberg discovery rate <0.05. Note that the compiled proteomic data is listed in Supplemental Table 1. (E) Total amount of metabolites detected in CAF-generated sEVs vs. DNPs, normalized to total EV protein levels. (F) Heat map depicting CAF sEV vs. DNPs normalized to median EV metabolite levels. Shown are the significant products using a t-test p-value of <0.1. A full readout of metabolomic analysis can be found in Supplemental Table 3.

### NetG1 and Int.α_5_ are enriched in sEVs collected from plasma of PDAC patients

Because CAFs comprise such a significant proportion of the cellular component in PDAC tumor masses, we posited that sEVs collected from human PDAC patient plasma could be enriched with CAF-generated vesicles. Hence, we questioned the levels of NetG1 and/or Int.α_5_ cargo in sEVs isolated from human PDAC patient plasma and compared these to levels detected sex and age matched healthy volunteers’ plasma. Excitingly, comparing 6 PDAC patient and 4 healthy control samples, we observed a 2.2 and 1.4 fold increase in NetG1 and Int.α_5_ levels, respectively, in PDAC patient plasma compared to levels detected in healthy volunteers (Figure 8A-C). Importantly, similarly to *in vitro* results, we did not detect significant differences in the amount of CD81 that was detected across all samples (Figure 8D). Together, these data provide evidence that NetG1 and Int.α_5_ could serve, in the future, as potential systemic biomarkers indicative of PDAC supportive stroma statuses in patients.

**Figure 8:**
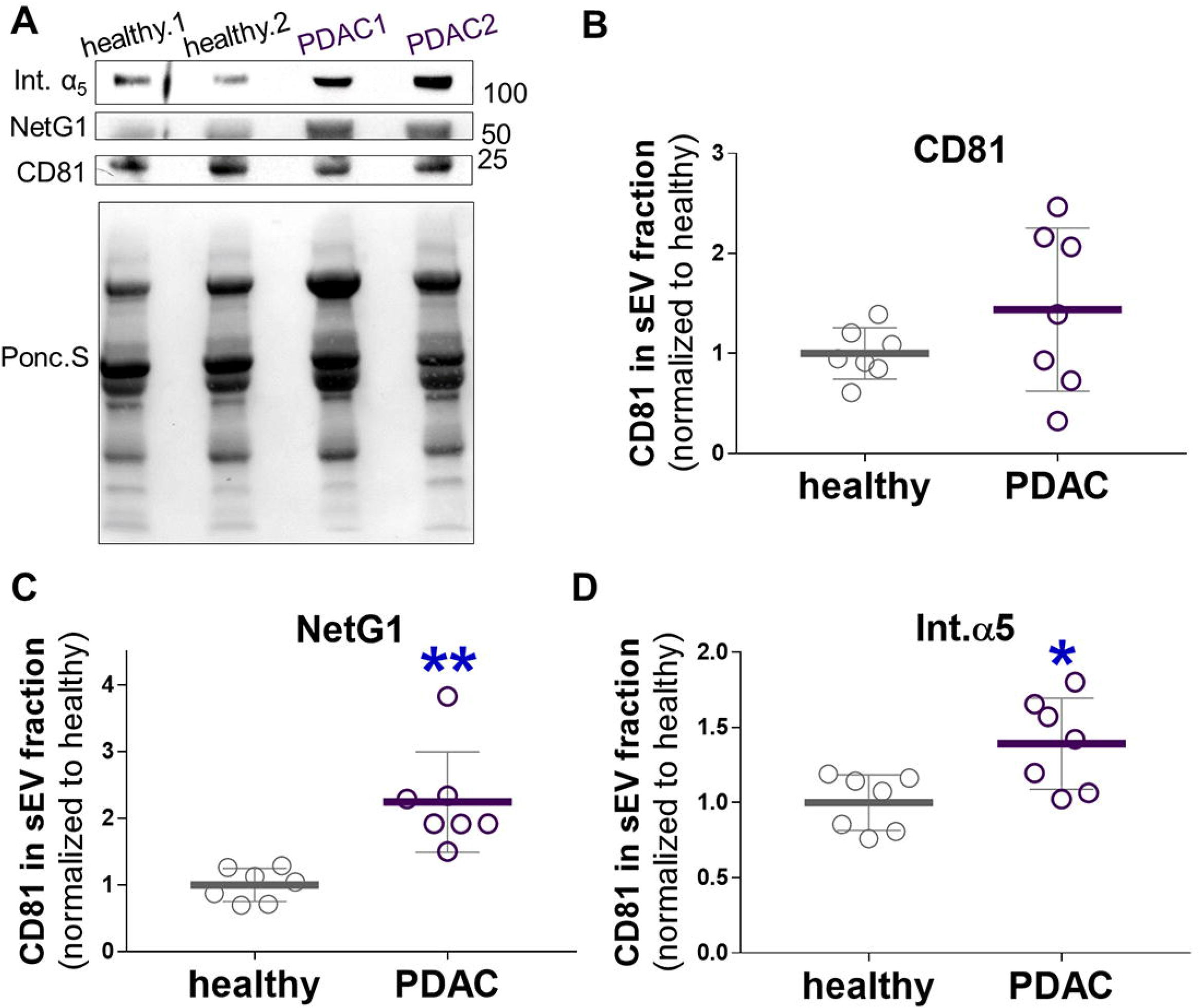
NetG1 and Int.α5 are enriched in sEVs collected from PDAC Patient Plasma. (A) Representative western blots of sEV fractions collected from plasma of PDAC patients (n =6) and healthy age/sex-matched control subjects (n=4). Lower panel depicts the Ponceau S stained of the sample and served as total protein loading control. (B-D) Quantification of CD81, NetG1, and Int.α_5_, levels, respectively, in sEVs loaded as in (A) and normalized to one arbitrary unit equaling the mean healthy volunteer (control) levels relative to each corresponding total protein amount gauged from the Ponceau S (A). Statistical test used: Unpaired t-test, with Welch’s correction. A comprehensive list of statistical readouts is provided in the Supplemental Table 2 file (Tabs = Fig.8B-D).

## Discussion

The PDAC tumor mass uniquely includes a significant stroma expansion of its desmoplastic TME with CAFs being the most common cell type. Four facts were considered in the central premise that was tested in this study: i) all cells are known to generate EVs ([46]); ii) tumor-promoting CAFs express NetG1 and encompass high levels of intracellular/endosomal activated Int.α_5_ that are most commonly detected in multivesicular bodies [25, 26], sub-organelles known to generate canonical exosomes ([47]); iii) tumor-promoting CAFs secrete factors that sustain PDAC cells from dying under nutrient-deprived conditions in a paracrine CAF NetG1-dependent manner [25]; and iv) there is reported evidence to suggest that CAF-generated EVs provide a metabolic benefit PDAC cells [15]. Hence, the hypothesis tested in this study was that NetG1 drives tumor-promoting CAFs’ ability to generate unique EVs that prevent PDAC apoptosis by providing a paracrine survival benefit to these cells, which are constantly stressed *in vivo* by limited nutrition.

sEV fractions are known to be enriched in exosomes and similarly sized vesicle structures, however this fraction is inherently heterogeneous ([48]). Through transmitted electron microscopy combined with double immuno-labeling, we detected NetG1 in exomere-like ([49]) structures that are much smaller than canonical exosomes. Notably, NetG1 failed to co-localize with known exosomal proteins like CD81 ([50]), yet the latter particles were co-labeled with anti-Int.α_5_ antibodies. Importantly, detecting Int.α_5_ in CAF-derived EVs in this study was in line with our previous work reporting that the CAF multivesicular cell body enrichment of activated Int.α_5_ is indicative of poor PDAC and renal cell carcinoma patient prognoses [26]. Regarding NetG1 localization at the PM, results from this study are in agreement with concurrent discoveries in the field which describe similar sub-exosomal structures as having a unique biogenesis pathway from exosomes and as a consequence, display distinct biochemical properties [21, 40]. Several studies have shown that DNP fractions are enriched in glycolytic components [20, 40], which served as a basis for using Hexokinase1 and GAPDH as positive markers for our DNP fraction and suggest a possible metabolic role for the newly identified NetG1^+^ vesicles.

While the exact mechanism of biogenesis and secretion of DNPs remains under investigation, previous studies have suggested that these vesicles can “shed off” from the PM [21], in a way that distinguishes these from the canonical pathway by which exosomes are generated (e.g., involving endosomal trafficking) ([51]). Of note, exomere-like DNPs have been shown to be enriched with GPI-anchored proteins ([49]). These studies could explain our observation that NetG1, also being a GPI-anchored protein [24], was detected in tumor-promoting CAFs at the PM, and was sorted to DNPs. Results from both the proteomic and metabolomic analyses also support the idea that EV heterogeneity renders specific functions to unique vesicles.

Regarding the proteomic signatures: both fractions included ECM-relevant components, which also supports previous work [52]. However, it is interesting to note that unique ECM proteins were segregated per fraction. For example, collagen alpha-2 was highly represented in DNPs while collagen alpha-1 and alpha-3 were enriched in sEVs. This observation is noteworthy as ratios of collagen chains in the ECM have been shown to inform pathological states in cancers [53]. Thus, further investigation into these trends may highlight specific EVs associated with stromal states. In addition, the CAF DNP fractions were enriched in proteins involved in cellular metabolism, which may not only serve as a way to positively identify these types of EVs but may also shed some light into how these DNPs exert a PDAC cell survival benefit. For example, DNPs included transketolase, a thiamine pyrophosphate-dependent enzyme involved in the pentose-phosphate metabolic pathway [54] and an integral component of the glucose-repurposing pathway for generating NADPH, nucleic acid, and amino acids [55]. Lactate Dehydrogenase A (LDHA), another key enzyme in glucose metabolism, was also highly represented in CAF generated DNPs. LDHA has been reported to be over expressed in cancer cells, and contributes to several oncogenic behaviors, the most relevant of which is maintaining cell survival under nutrient-stress [56].

While both sEVs and DNPs performed a function in supporting the survival of nutrient deprived PDAC cells, the mechanisms that drive each may differ. Metabolomic data demonstrated that CAF sEVs have a substantial NetG1-dependent metabolite profile, likely required for pro-survival behavior. Notably, sEVs have significantly more enriched metabolite content compared to DNPs, suggesting that the pro-survival function observed in DNPs could be due to their protein content instead. Importantly, previous studies have demonstrated the ability of EVs to transfer functional enzymes to recipient cells [17, 40, 57, 58], thus there is evidence supporting DNP-mediated function could be attributed to the transfer of metabolism- and other stress-response related enzymes; while sEV-mediated survival could stem from metabolite content. In an *in vivo* context, the EV-mediated transfer of these types of metabolism-stimulating proteins and key metabolites, such as glutamine, from CAFs to PDAC could be a mechanism by which CAFs are able to provide pro-survival advantages during early disease development. Data collected in this study regarding the specific metabolites packed as NetG1^+^ CAF-generated sEV cargo, nicely matched our previous results reporting that CAFs modulate pro-tumoral metabolism via two key proteins [25]. The first was glutamine synthetase, known for the synthesis of glutamine and the second was VGLUT1, which is a well-characterized glutamatergic transporter responsible for loading presynaptic vesicles. Both proteins were shown to be modulated by and act downstream to NetG1 in PDAC CAFs and both were reported to be necessary for NetG1 pro-tumoral function [25]. Ultimately, it will be the goal of future studies to further elucidate the NetG1-modulated mechanisms responsible for sorting cargo and generating the novel EV subpopulations produced by NetG1^+^ CAFs.

We also note that losing NGL1 in PDAC cells eliminated the benefit provided by CAF EVs. Our previous work showed that NGL1-deficiency in these cells causes a decrease in the amount of macropinocytosis mediated uptake of extracellular material, which can be one potential explanation for the loss of function in survival following CAF sEV treatment. Interestingly, this function was reported to be independent of NetG1 [25]. Then again, a NetG1/NGL1 dependent material transfer was also reported [25], which was too evident in this study, based on results obtained using increased amounts of rNGL1. These results could be due to the rNGL1 engaging NetG1 on the EVs, and thus preventing an interaction with NGL1 on the PDAC cells, and blocking subsequent uptake of EVs. Considered as a whole, there is reported evidence for a NetG1-independent mechanism of CAF EV-mediated survival regulated by macropinocytosis, as well as a NetG1-NGL1 dependent mechanism further supported by results from this study.

One exciting aspect of this study is the fact that we found evidence to suggest that the unique EVs we herein report, identified *in vitro* using human CAFs cultured under pathophysiologic conditions, can also be detected in plasma from PDAC patients and levels of these unique vesicles are significantly higher than levels in plasma from healthy donors. This is important when considering that PDAC is a difficult disease to detect systemically. The development of a blood-based PDAC-associated EV profile would serve a much-needed role in the early detection of this disease [59]. Hence, our findings establish a basis for a future in-depth study that could query the potential biomarker value of unique EVs as indicators of desmoplastic/stromal states. To this end, numerous recent studies have demonstrated that stroma signatures could be indicative of PDAC and other cancers’ patient outcomes [25, 60–68] .

EV research is a broad yet rapidly expanding field that continues to elucidate the diverse nature of EVs both from a functional, as well as a characteristic standpoint. In this study, we report the novel role of NetG1 in CAF-EV mediated support of nutrient-deprived PDAC cells, as well as early evidence to their potential usefulness as clinical biomarkers.

## Methods

### Statement of Ethics

The authors state the work contained in this study was conducted in accordance with accepted ethical guidelines. All facilities within Fox Chase Cancer Center adhere to the International Society for Biological and Environmental Repositories and National Cancer Institute Best Practices for Biospecimen Resources. All human samples used were donated with informed consent for the purpose of research and all were decoded for research usage.

### Key Resources Table

All key resources used in this study are listed in the “Key Resources Table”, provided as a supplemental file. The sources and identifier codes are provided therein.

### Parental cell lines

Patient matched CAFs (and selected tumor adjacent fibroblasts) were collected from fresh surgical tissue and enzymatically digested and characterized using our well-established protocols [31, 33]. PANC-1 and AsPC-1 were obtained from ATCC. HPNE and isogenic KRAS-HPNE (HPNE cells with E6/E7-KRASG12D mutations) cells [36], were also from ATCC. All experimental data includes cells that were used within 3-15 passages after being thawed. Fibroblastic cells were authenticated for species of origin and reported genetic profiles [25, 26]. All cells were tested regularly for mycoplasma contamination using PCR detection.

### Engineered cell lines

Engineered cell lines used in this study were generated and validated as published in previous works include immortalized CAF and NLF (generated by knocking down the beta5-integrin subunit [26]) cell lines, Ctl. and NetG1-KD CAF lines, mCherry-expressing PDAC cell lines, and NGL-1 KD PANC-1 cells [25, 26, 31]. Cells engineered specifically for this project include parental CAFs (Ctl. and NetG1-KD) overexpressing green fluorescent protein (GFP). Briefly, eGFP was PCR amplified using Phusion HF polymerase reagent master mix, then cloned into Xbal/Xhol digested Fast AP dephosphorylated pLV-CMV-H4-puro vector Ctl. and NetG1-KD CAFs were then transduced with pLV-CMV-H4-puro-GFP with 10 μg/mL polybrene. Cells were selected in media containing 2 μg/mL puromycin and GFP-expressing cells were selected using flow cytometry. Parental NetG1-KD CAF cells overexpressing (OE) ectopic NetG1 (NetG1-OE) were generated as follows: NetG1 mRNA was isolated from CAFs and converted to cDNA using the SuperScript™ IV Reverse Transcriptase kit (Thermo Fisher Scientific). NetG1 cDNA was PCR amplified using primers that contained XbaI/XhoI overhangs, and the PCR product was cloned into the XbaI/XboI cut (New England Biolabs) and dephosphorylated (Fast AP, Thermo Fisher Scientific) pLV-CMV-H4-puro overexpression vector.

### Cell Culture and Condition Media Collection

All fibroblasts were cultured in DMEM containing 4.5g/L glucose, 1.5g/L NaHCO_3_, 10% Fetal Bovine Serum, 4mM L-glutamine (Corning, Corning, NY), and 1% penicillin/streptomycin (Invitrogen, Waltham, MA). The methodology of our 7-day 3D-cell derived ECM (also recognized as CDM) production has been published previously [33]. At the completion of 3D matrix production, cells were rinsed twice with PBS and switched to a serum/EV-free DMEM (1%pen/strep, 4 μM glutamine) and were sustained for 48 hours at 37°C, to generate conditioned media (CM). Upon completion of the conditioning period, CM was removed to be processed for further experiments, and the cells and CDMs were lysed for protein extraction (see methods below for depiction of metabolite extraction). All PDAC cell lines used in this study were maintained in 4:1 DMEM (1g/mL glucose, 110 mg/mL sodium pyruvate) M3 Base (INCELL, San Antonio, TX) supplemented with 5% FBS, and 1% penicillin-streptomycin, until needed for experimental purposes.

### ECM Immunofluorescent Labeling and Image Acquisition

Fibroblast CDMs were imaged using immunofluorescent labeling as described [31]. Cells and ECMs were immunolabeled using primary antibodies against Fibronectin (rabbit IgG; 92.4 μg/mL) and α-smooth muscle actin (αSMA) (mouse IgG; 75) for 1 hour at RT. Following primary immunolabeling, CDM cultures were washed with a solution of PBS and 0.05% TWEEN (PBST) 3 times for 5 minutes each. Biomarkers were labeled using host-specific secondary antibodies containing conjugated fluorophores for 1 hour at RT, so that fibronectin could be visualized at 647nm, and αSMA at 568nm. Following secondary antibody incubations, the CDMs were washed as before. SYBR Green (Invitrogen, Waltham, MA), a DNA stain that emits at 520nm, was added to the cultures at [1:10,000] for 20 minutes at RT to visualize nuclei. Following SYBR Green treatment, CDM cultures were again rinsed and mounted onto a glass microscopy slides with Prolong™ Gold Anti-fade Mountant (Invitrogen, Waltham, MA). Samples were stored at +4°C for 48-96 hours to allow the mounting medium to set, before visualizing using spinning disk confocal microscopy (Perkin Elmer, Waltham, MA) with a Nikon Eclipse T*i*2 inverted microscope at 60x. Excitation-emission spectra used for the following channels: FITC (488 nm–520 nm), TRITC (540 nm–580 nm), CY5 (640–670 nm). Images were acquired on a Hamamatsu Orca-flash 4.0 camera (Hamamatsu, Japan), taking 6-8 images per coverslip covering a representative area of the entire coverslip. The top and bottom of each section being recorded was used to create a z-stack for each fluorescent channel. Images were then exported and processed via FIJI [69] for analysis as described in [31].

### Media fractionation; sEV Isolation and preparation of functional conditioned media

sEVs were purified from conditioned media using differential centrifugation [42]. Briefly, conditioned media, collected as described above, was centrifuged for 10 minutes at 300 x *g*, followed by a 20-minute centrifugation at 2000 x *g* to pellet cellular debris and dead cells. The resulting supernatant was spun for 30 minutes at 12,000 x *g* to pellet large vesicles and free-floating organelles. The new supernatant was filtered through a 0.22 μm PDVF membrane using a syringe pump. The filtered sample was then either used as “functional conditioned media” for survival assays or transferred to a Beckman Optima TL-100 Ultracentrifuge (Beckman Coulter Life Sciences, Brea, CA) for sEV isolation. To isolate sEVs, the filtered sample was spun down at 120,000 x *g* for 2 hours to pellet sEVs. The sEV supernatant was either used as an experimental condition or stored at +4°C to later isolate DNPs (see below). The pellet containing sEV was washed with 1mL PBS and spun again at 120,000 x *g* for 2 hours to concentrate the pellet of sEVs. The resulting sEV pellet was re-suspended in 50 μL PBS, and either used immediately for functional assays, or stored at −80°C for added analytical characterization.

### Isolation of DNPs

Distinct nanoparticle fractions (DNPs) were enriched from the supernatant of the sEV fraction (see above) by an additional ultracentrifugation stage that was adapted from Zhang *et. al*. [40]. The sample was spun at 120,000 x *g* for 16 hours to pellet DNPs (a sub-exosomal subset of extracellular vesicles). The DNP-containing pellet was washed with 1 mL PBS and spun again at 120,000 x *g* for 2 additional hours. The resulting DNP pellet was re-suspended in 50 μL PBS, and either used immediately for functional assays, or stored at −80°C for added analytical characterization.

### Human Plasma Acquisition and sEV isolation

A total of ten human plasma samples (6 PDAC patients and 4 healthy donors) were used in this study. All individuals were white male of about 60 years of age. Samples were collected through Fox Chase Cancer Center’s Institutional Biosample Repository Facility following a signed informed consent, using HIPAA approved protocols and exemption-approval of Fox Chase Cancer Center’s Institutional Review board. Subjects’ identities were protected by classified coded identification numbers. Collected blood samples were processed to isolate plasma, and stored at −80°C. Once thawed, sEV isolation from plasma was performed similarly to that from cell-conditioned media with the following modifications: Plasma was diluted 1:1 with PBS, then centrifuged for 10 minutes at 500 x *g*, supernatant was collected spun for 30 minutes at 2000 x *g*. The resulting supernatant was spun for 45 minutes at 14,000 x *g*. This supernatant was filtered through a 0.22μm PDVF membrane using a syringe pump. The filtered sample was spun down at 150,000 x *g* for 2 hours to pellet sEVs. The sEV pellet was washed with 1mL PBS and spun again at 150,000 x *g* for 2 hours to concentrate the pellet of sEVs. The sEV pellet was re-suspended in 150 μL PBS and stored at −80°C.

### Nanoparticle quantification

Fractionated EV compositions were analyzed using two machines and their respective proprietary software: the ZetaView PMX-120 (ParticleMetrix, Germany), and the NanoSight NS300 (Malvern Pananalytical, Malvern PA). Isolated EV samples were diluted 1:100 in sterile PBS (twice filtered through a 0.22μm PDVF membrane) to a final volume of 1mL. The diluted sample was then aspirated into a sterile 1mL syringe and pumped into the Nanosight machine. Three 30-second videos were recorded wherein the sample was advanced progressively between each recording. These recorded videos were processed by the respective software, to produce a readout including the total particle count of the sample, in addition to batching the particle sizes with the concentration of particles for each size range.

### Cell Viability Assay

mCherry-expressing PDAC cells were rinsed 2x with PBS and switched to low glucose (1.5 g/mL), serum- and glutamine-free DMEM (1.5g/mL glucose) for 6 hours to subject cells to pathophysiologic-like nutritional stress, prior to seeding in a 96-well clear-bottom black microplate (Grenier, Germany) at 7.5×10^3^ cells/well in rows B-G; cells were incubated at 37°C overnight. The following day, cell media was removed, and treatment conditions were administered to cells as described in figure legends. Separate conditions were arranged by column, providing 6 technical replicates with acellular wells at the top(A) and bottom(H) to act as media blanks per condition. Cell were incubated at 37°C for 48 hours post-treatment (unless stated otherwise) and gauged for cell viability. We defined “cell viability” as the ratio of live cells to dead cells. Live cells were measured via mCherry expression. Cell death was measured via treatment with Sytox Blue (Thermo Fisher Scientific, Waltham, MA). Sytox Blue was diluted in PBS and added to the media of all wells at a final concentration of 0.5 μM and incubated in the dark at 37°C for 15 minutes, according to the manufacturer’s protocol. After Sytox treatment, fluorescence was measured using a Tecan Spark™ 10M microplate reader (Tecan, Switzerland). Live cells/mCherry expression were recorded using 540 nm (excitation)/590 nm (emission), and Cell Death/sytox using 430 nm (excitation)/485 nm (emission); which was measured in that order to minimize residual autofluorescence between channels. Z-position and gain values were optimized to the positive control condition. Raw data for each fluorescence channel was processed by averaging the 2 blank values for each condition and subtracting the averaged blank value from the values of their respective conditions’ technical replicates. Blank-adjusted values for each channel were then used to divide the live cell value by the death cell value for each matched well. The cell viability ratios for each well were then normalized to the average of all control (PBS/untreated) values to identify each condition’s fold change in cell viability compared to the control.

### PDAC/Fibroblast Direct Co-culture

mCherry-expressing PDAC cells were switched to low glucose (1.5 g/mL), serum/glutamine free media for 24 hours and seeded at 7.5*10^3^ into a 96-well format containing fully confluent wells of pre-seeded fibroblasts and their secreted CDMs and incubated at 37°C for an additional 48 hours. After 48 hours, the plate was washed with PBS to remove dead cells and debris, before being analyzed on a Tecan Spark™ 10M microplate reader (Tecan, Switzerland). Numbers of PDAC cells were measured by quantifying mCherry expression attained using 540 nm (excitation)/590 nm (emission). PDAC levels were normalized to the average of all single-cultured PDAC cell values to identify each condition’s fold change in PDAC cell count.

### Transmission Electron Microscopy (TEM)

EVs were fixed in 2% Paraformaldehyde (Electron Microscopy Sciences, Hatfield, PA) and 20 μL of fixed EV suspension was carefully placed on top of a Formvar-coated Film, 300-mesh nickel grid (Electron Microscopy Sciences, Hatfield, PA). To attain contrast stain, the grid was incubated onto a 30 μL droplet of PBS, twice, for 5 minutes, and on a 30 μL droplet of 2% uranyl acetate solution (Electron Microscopy Sciences, Hatfield, PA) for 10 additional minutes at RT, Excess Uranyl acetate was removed and the gird was stored in the dark at RT for future use. For immune-labeling, the grid was placed on a 30 μL droplet of Odyssey Blocking Buffer; PBS-base (Li-Cor, Lincoln, NE) for 10 minutes at RT. The sample was then transferred onto a 30 μL droplet of primary antibody diluted in blocking buffer for 1 hour at RT. Antibodies used: CD81 [1 μg/mL], NetG1 [1 μg/mL], SNAKA51 [70] [0.225 μg/mL]. Samples were rinsed 3 times with PBS for 5 minutes at RT prior to transferring onto a 30 μL droplet of corresponding secondary antibodies conjugated to either a unique sized gold particle or a unique quantum dot, diluted in blocking buffer, for 1 hour at RT. Samples were rinsed as before and incubated 2% uranyl acetate solution for 10 minutes as above. EVs were visualized under an JEOL 2100 transmission electron microscope (JEOL, Ltd., Japan) at 60,000x-135,000x magnification.

### SDS-PAGE and Western Blot Analysis

Whole cell lysates were prepared for Western Blotting by lysing 10^6^ cells in a 6-well format, using 100-200 μL of *radio-immunoprecipitation assay* (RIPA) buffer as previously made [25]. Protein concentrations of samples were determined using a Bradford assay [71] and working samples were normalized to an equal concentration (approximating 5 μg protein loaded per sample). To immunodetect protein, the following primary antibodies were used: Int.α_5_ [0.1 μg/mL], Alix [0.2 μg/mL], αSMA, [7.5 μg/mL], CD63 [0.2 μg/mL], CD81 [0.067 μg/mL], GAPDH [0.05 μg/mL], GFP [0.014 μg/mL], Histone 3 [0.0006 μg/mL], Hexokinase 1 [0.8 μg/mL], NetrinG1 Ligand [1.0 μg/mL], NetG1 [1.6 μg/mL], PARP [0.39 μg/mL], TSG101 [0.44 μg/mL]. secondary antibodies used were as follows: anti-mouse HRP [0.03 μg/mL], anti-rabbit HRP [0.013 μg/mL]. Immunoblots were visualized by treating membranes with Immobilon Western Chemiluminescent HRP substrate (Millipore, Burlington, MA), exposed, and developed on autoradiography film.

### sEV-mediated transfer of GFP

sEVs were isolated from GFP-expressing CAFs, and then administered to PDAC cells to assess GFP transfer via Western Blot. AsPC-1 cells were rinsed 2x with PBS and switched to low glucose (1.5 g/mL), serum/glutamine free DMEM for 6 hours, before being plating at 10^3^ cells/well in a 6-well format. These cells were incubated for 24 hours in serum/glutamine free media, before being rinsed 1x with PBS and treated with equal amounts of sEVs suspended in 2mL PBS from GFP-expressing Ctl. or NetG1-KD CAFs. Equal volumes of sEVs from each condition were stored at −80°C to be used as input controls for Western Blot analysis.. 24 hours post-treatment, supernatant was removed, cells were rinsed 2x and lysed with RIPA buffer and processed for Western Blot analysis.

### Proteomic Sample Processing, Mass Spectrometry, and Spectra Analysis

Approximately 50μg of EV proteins were reduced with 10 mM dithiothreitol for 25 min, alkylated with 50 mM indole-3-acetic acid for 30 minutes in the dark, and precipitated overnight with 80% acetone. The precipitated pellet was washed once with 80% acetone, re-suspended in 6 M urea/2 M and digested using LysC and trypsin. Digested peptides were cleaned using stage tips, eluted with 70% acetonitrile and lyophilized. Samples were re-suspended in 0.1% formic acid and separated with a Thermo Scientific RSLCnano Ultimate 3000 LC on a Thermo Scientific Easy-Spray C18 PepMap 75μm x 50cm C-18 2 μm column. A 75 min gradient of 2-25% (30 min) acetonitrile with 0.1% formic acid was run at 300 nL/min at 50°C. Eluted peptides were analyzed by a Thermo Scientific Q Exactive mass spectrometer utilizing a top 15 methodology in which the 15 most intense peptide precursor ions were subjected to fragmentation. The automatic gain control for MS1 was set to 3×106 with a max injection time of 120 ms, the automatic gain control for MS2 ions was set to 1×105 with a max injection time of 150 ms, and the dynamic exclusion was set to 90 s.

### Proteomics Data Processing

Raw data analysis of label free quantitation (LFQ) experiments was performed using MaxQuant software 1.6.1.0[72] and searched using Andromeda 1.5.6.0 [73] against the Swiss-Prot human protein database (downloaded on April 24, 2019, 20402 entries). The search was set up for full tryptic peptides with a maximum of two missed cleavage sites. All settings were default and searched using acetylation of protein N-terminus and oxidized methionine as variable modifications. Carbamidomethylation of cysteine was set as fixed modification. The precursor mass tolerance threshold was set at 10 ppm and maximum fragment mass error was 0.02 Da. LFQ quantitation was performed with the following parameters: LFQ minimum ratio count: 1 global parameters for protein quantitation were as follows: label minimum ratio count: 1, peptides used for quantitation: unique, only use modified proteins selected and with normalized average ratio estimation selected. Match between runs was employed for LFQ quantitation and the significance threshold of the ion score was calculated based on a false discovery rate of < 1%. MaxQuant normalized LFQ values were imported into Perseus software (1.6.2.3) [74] and filtered in the following manner: kinases identified by site only were removed, reverse, or potential contaminants were removed. Protein LFQ values were log2 transformed and subjected to the Student’s t-test comparing sEV and DNP fractions. Parameters for the Student’s t-test were the following: S0=2, side both using Benjamini-Hochberg false discovery rate (FDR) <0.05.

### Protein Gene Ontology Analysis

Gene Ontology was performed using Metascape.org’s open access gene annotation and analysis resources [30] following proteomic data processing. Statistically significant protein signatures identified using the Perseus software were entered into Metascape and run using their “express analysis” function. Using this list of proteins, Metascape’s software identifies all statistically enriched terms including GO/KEGG terms, canonical pathways, and hallmark gene sets. Accumulative hypergeometric p-values and enrichment factors were also calculated and used to filter results, which were clustered into a hierarchical tree based on Kappa-statistical similarities in gene membership. Then a 0.3 kappa score was applied as the threshold to cast the tree into term clusters ranked by p-value. Enrichment criteria including a minimum overlap value of 3, minimum enrichment value of 1.5, and a p-value cutoff of 0.01 were applied. A full report for each experiment is included in supplementary data tables labeled “Enrichment Analysis”.

### sEV boiling/filtering

Crude metabolite contents of sEV and sEV supernatant fractions were collected by boiling samples. sEV and supernatant fractions were isolated as described above. Samples were resuspended in 500 μL sterile PBS and incubated at 100°C for 10 minutes. Samples were allowed to return to RT before being filtered through Amicon Ultra centrifuge filters, 3000kDa pore size (Millipore, Wilmington, DE) for 30 minutes at 14,000*g*. Equal volumes of filtrate volume were collected, brought to a total volume of 900 μL sterile PBS and used in cell viability assays as described.

### Metabolite sample collection

CAF sEV and DNP fractions were isolated from conditioned media as described above. sEVs were resuspended in 200 μL PBS and 50 μL aliquots were taken for BCA protein quantification. The remaining volume was incubated with ice cold methanol (80% by volume) for 10 min, inverting over dry ice. Sample was centrifuged at 17,000g for 10 min and supernatant was transferred to a clean tube. Metabolite supernatants were normalized to total protein content, and normalized samples were desiccated in a speed vacuum centrifuge and stored at −80oC until being processed. When ready, samples were suspended in a 50:50 mixture of methanol and water in HPLC vials for LC-MS/MS analysis and were run using the following protocol.

### Snapshot metabolomics

Samples were run on an Agilent Technologies 1290 Infinity II LC-6470 Triple Quadrupole (QqQ) tandem mass spectrometer (MS/MS) system with the following parameters: 1290 Infinity II LC Flexible Pump (Quaternary Pump), 1290 Infinity II Multisampler, 1290 Infinity II Multicolumn Thermostat with 6 port valve and 6470 triple quad mass spectrometer. Agilent Masshunter Workstation Software LC/MS Data Acquisition for 6400 Series Triple Quadrupole MS with Version B.08.02 is used for compound optimization and sample data acquisition.

Solvent A is 97% water and 3% methanol 15 mM acetic acid and 10 mM tributylamine at pH of 5. Solvent C is 15 mM acetic acid and 10 mM tributylamine in methanol. Washing Solvent D is acetonitrile. LC system seal washing solvent 90% water and 10% isopropanol, needle wash solvent 75% methanol, 25% water. The materials used for the solvents were: GC-grade Tributylamine 99% (ACROS ORGANICS), LC/MS grade acetic acid Optima (Fisher Chemical), InfinityLab Deactivator additive, ESI –L Low concentration Tuning mix (Agilent Technologies), LC-MS grade solvents of water, and acetonritele, methanol (Millipore), isopropanol (Fisher Chemical).

An Agilent ZORBAX RRHD Extend-C18, 2.1 × 150 mm, 1.8 um and ZORBAX Extend Fast Guards for UHPLC are used in the separation. LC gradient profile is: at 0.25 ml/min, 0-2.5 min, 100% A; 7.5 min, 80% A and 20% C; 13 min 55% A and 45% C; 20 min, 1% A and 99% C; 24 min, 1% A and 99% C; 24.05 min, 1% A and 99% D; 27 min, 1% A and 99% D; at 0.8 ml/min, 27.5-31.35 min, 1% A and 99% D; at 0.6 ml/min, 31.50 min, 1% A and 99% D; at 0.4 ml/min, 32.25-39.9 min, 100% A; at 0.25 ml/min, 40 min, 100% A. Column temp is kept at 35 □C, samples are at 4 □C, injection volume is 2 μl.

6470 Triple Quad MS is calibrated with the ESI-L Low concentration Tuning mix. The source parameters are: Gas temp 150 □C, Gas flow 10 l/min, Nebulizer 45 psi, Sheath gas temp 325 □C, Sheath gas flow 12 l/min, Capillary −2000 V, Delta EMV −200 V. Dynamic MRM scan type is used with 0.07 min peak width and the acquisition time is 24 min. dMRM transitions and other parameters for each compound are listed in a separate sheet. Delta retention time of plus and minus 1 min, fragmentor of 40 eV and cell accelerator of 5 eV are incorporated in the method.

### Metabolomics Data Analysis

The QqQ data were pre-processed with Agilent MassHunter Workstation QqQ Quantitative Analysis Software (B0700). Each metabolite abundance level in each sample was divided by the median of all abundance levels across all samples for proper comparisons, statistical analyses, and visualizations among metabolites. The statistical significance test was done by a two-tailed t-test with a significance threshold level of 0.10. All error bars represent mean with standard deviation. Heatmaps were generated and data clustered using Morpheus by Broad Institute (https://software.broadinstitute.org/morpheus). Pathway analyses were conducted using MetaboAnalyst (https://www.metaboanalyst.ca).

### Statistics

Prism 7.05 (Graph Pad Software, San Diego, California) was used for all statistical analysis. For comparison between two groups, a two-tailed unpaired t-test was performed. For comparisons between more than two groups, a One-Way ANOVA was performed, using either a Dunnett’s multiple comparisons test (compared to control condition) or Tukey’s multiple comparisons test (comparing all conditions to one another), unless otherwise noted. Groups were deemed statistically significantly different from one another if the p-value was smaller than or equal to 0.05. On graphs, the symbols “*” and “#” were used to denote significance and defined as follows: *(#) p<0.05; **(##) p<0.01; ***(###) p< 0.001; ****(###) p< 0.0001. Comprehensive statistics readouts provided in in the excel file labeled Supplemental Tables 2.

## Supporting information

Key resources table

## Acknowledgments

We dedicate this work to Patricia Keely (pioneer in the field of ECM biology) and Neelima Shah (the most beautiful soul) who continue to inspire our studies. We thank Jane Clifford, Paul Campbell, and Jeffery Peterson for assertive comments during the time experiments were being conducted. We are grateful to Martin Humphries for the SNAKA51 antibody. We thank Rachel DeRita for her insight and expertise in analyzing extracellular vesicles. This work was supported in part by the Pancreatic Cancer Cure Foundation, the Concetta Greenberg Pancreatic cancer Institute, Pennsylvania’s DOH Health Research Formula Funds, the 5th AHEPA Cancer Research Foundation, Inc., a grant by the Worldwide Cancer Research, NIH/NCI grants R21 CA231252, R21CA252535, R01CA232256, and the Core Comprehensive Cancer Center Grant CA06927 in support to Fox Chase Cancer Center’s facilities including: Bio Sample Repository, Light Microscopy, Biostatistics and Bioinformatics, Immune Monitoring, Cell Culture, and the Talbot Library. The authors also acknowledge grants from the ACS-132561-PF-18-218-01-CSM (JCG), the 2021 Pancreatic Cancer Action Network Career Development Award in memory of Skip Viragh 21-20-FRAN (RF).

## Author contributions

Conceptualization and writing (KSR and EC); data analyses and/or significant experimental contributions (KSR, RF, JFB, JG, DBVC, TL, NP, AA, AK, CO, JSD, and CAL), discussions with significant contributions (KSR, RF, DBVC, CAL, LRL and EC), final text editing and concepts (all authors).

## References

1. Rawla, P., T. Sunkara, and V. Gaduputi, Epidemiology of Pancreatic Cancer: Global Trends, Etiology and Risk Factors. World J Oncol, 2019. 10(1): p. 10–27.

2. SEER*Stat Database Incidence - SEER Research Data, R., Nov 2020 Sub (1975-2018) - Linked To County Attributes - Time Dependent (1990-2018) Income/Rurality, 1969-2019 Counties, National Cancer Institute, DCCPS, Surveillance Research Program, released April 2021, based on the November 2020 submission., 2019.

3. Siegel, R.L., et al., Cancer Statistics, 2021. CA Cancer J Clin, 2021. 71(1): p. 7–33.

4. Rahib, L., et al., Estimated Projection of US Cancer Incidence and Death to 2040. JAMA Network Open, 2021. 4(4): p. e214708–e214708.

5. Mahadevan, D. and D.D. Von Hoff, Tumor-stroma interactions in pancreatic ductal adenocarcinoma. Mol Cancer Ther, 2007. 6(4): p. 1186–1197.

6. Piersma, B., M.K. Hayward, and V.M. Weaver, Fibrosis and cancer: A strained relationship. Biochim Biophys Acta Rev Cancer, 2020. 1873(2): p. 188356.

7. Drifka, C.R., et al., Highly aligned stromal collagen is a negative prognostic factor following pancreatic ductal adenocarcinoma resection. Oncotarget, 2016. 7(46): p. 76197–76213.

8. Kerk, S.A., et al., Metabolic networks in mutant KRAS-driven tumours: tissue specificities and the microenvironment. Nat Rev Cancer, 2021. 21(8): p. 510–525.

9. Lyssiotis, C.A. and A.C. Kimmelman, Metabolic Interactions in the Tumor Microenvironment. Trends Cell Biol, 2017. 27(11): p. 863–875.

10. Son, J., et al., Glutamine supports pancreatic cancer growth through a KRAS-regulated metabolic pathway. Nature, 2013. 496(7443): p. 101–105.

11. von Ahrens, D., et al., The role of stromal cancer-associated fibroblasts in pancreatic cancer. J Hematol Oncol, 2017. 10(1): p. 76.

12. Doyle, L.M. and M.Z. Wang, Overview of Extracellular Vesicles, Their Origin, Composition, Purpose, and Methods for Exosome Isolation and Analysis. Cells, 2019. 8(7).

13. Bebelman, M.P., et al., Biogenesis and function of extracellular vesicles in cancer. Pharmacol Ther, 2018. 188: p. 1–11.

14. Yáñez-Mó, M., et al., Biological properties of extracellular vesicles and their physiological functions. J Extracell Vesicles, 2015. 4: p. 27066.

15. Zhao, H., et al., Tumor microenvironment derived exosomes pleiotropically modulate cancer cell metabolism. Elife, 2016. 5.

16. Mathivanan, S. and R.J. Simpson, ExoCarta: A compendium of exosomal proteins and RNA. Proteomics, 2009. 9(21): p. 4997–5000.

17. Singh, A., et al., Exosome-mediated Transfer of αvβ3 Integrin from Tumorigenic to Nontumorigenic Cells Promotes a Migratory Phenotype. Mol Cancer Res, 2016. 14(11): p. 1136–1146.

18. Harada, Y., et al., Extracellular Vesicles and Glycosylation. Adv Exp Med Biol, 2021. 1325: p. 137–149.

19. Cerezo-Magaña, M., A. Bång-Rudenstam, and M. Belting, The pleiotropic role of proteoglycans in extracellular vesicle mediated communication in the tumor microenvironment. Seminars in Cancer Biology, 2020. 62: p. 99–107.

20. Zhang, H. and D. Lyden, Asymmetric-flow field-flow fractionation technology for exomere and small extracellular vesicle separation and characterization. Nature Protocols, 2019. 14(4): p. 1027–1053.

21. Zhang, H., et al., Identification of distinct nanoparticles and subsets of extracellular vesicles by asymmetric flow field-flow fractionation. Nat Cell Biol, 2018. 20(3): p. 332–343.

22. Hoshino, A., et al., Extracellular Vesicle and Particle Biomarkers Define Multiple Human Cancers. Cell, 2020. 182(4): p. 1044–1061.e18.

23. Dun, X.P. and D.B. Parkinson, Role of Netrin-1 Signaling in Nerve Regeneration. Int J Mol Sci, 2017. 18(3).

24. Nakashiba, T., et al., Netrin-G1: a novel glycosyl phosphatidylinositol-linked mammalian netrin that is functionally divergent from classical netrins. J Neurosci, 2000. 20(17): p. 6540–50.

25. Francescone, R., et al., Netrin G1 promotes pancreatic tumorigenesis through cancer associated fibroblast-driven nutritional support and immunosuppression. Cancer Discovery, 2021. 11(2): p. 446–479.

26. Franco-Barraza, J., et al., Matrix-regulated integrin αvβ5 maintains α5β1-dependent desmoplastic traits prognostic of neoplastic recurrence. eLife, 2017. 6: p. e20600.

27. Latifkar, A., R.A. Cerione, and M.A. Antonyak, Probing the mechanisms of extracellular vesicle biogenesis and function in cancer. Biochem Soc Trans, 2018. 46(5): p. 1137–1146.

28. Zhang, X., et al., Exosomes in cancer: small particle, big player. J Hematol Oncol, 2015. 8: p. 83.

29. Théry, C., et al., Minimal information for studies of extracellular vesicles 2018 (MISEV2018): a position statement of the International Society for Extracellular Vesicles and update of the MISEV2014 guidelines. J Extracell Vesicles, 2018. 7(1): p. 1535750.

30. Zhou, Y., et al., Metascape provides a biologist-oriented resource for the analysis of systems-level datasets. Nat Commun, 2019. 10(1): p. 1523.

31. Franco-Barraza, J., et al., Engineering clinically-relevant human fibroblastic cell-derived extracellular matrices. Methods Cell Biol, 2020. 156: p. 109–160.

32. Hinz, B., et al., Alpha-smooth muscle actin expression upregulates fibroblast contractile activity. Molecular biology of the cell, 2001. 12(9): p. 2730–2741.

33. Franco-Barraza, J., et al., Preparation of extracellular matrices produced by cultured and primary fibroblasts. Curr Protoc Cell Biol, 2016. Chapter 10(71): p. 10.9.1–10.9.34.

34. Seiradake, E., et al., Structural basis for cell surface patterning through NetrinG-NGL interactions. EMBO J, 2011. 30(21): p. 4479–88.

35. Moore, S.W., M. Tessier-Lavigne, and T.E. Kennedy, Netrins and their receptors. Adv Exp Med Biol, 2007. 621: p. 17–31.

36. Campbell, P.M., et al., K-Ras promotes growth transformation and invasion of immortalized human pancreatic cells by Raf and phosphatidylinositol 3-kinase signaling. Cancer Res, 2007. 67(5): p. 2098–106.

37. Mizenko, R.R., et al., Tetraspanins are unevenly distributed across single extracellular vesicles and bias sensitivity to multiplexed cancer biomarkers. J Nanobiotechnology, 2021. 19(1): p. 250.

38. Kowal, J., et al., Proteomic comparison defines novel markers to characterize heterogeneous populations of extracellular vesicle subtypes. Proceedings of the National Academy of Sciences, 2016. 113(8): p. E968–E977.

39. Zhang, H. and D. Lyden, Asymmetric-flow field-flow fractionation technology for exomere and small extracellular vesicle separation and characterization. Nat Protoc, 2019. 14(4): p. 1027–1053.

40. Zhang, Q., et al., Transfer of Functional Cargo in Exomeres. Cell Rep, 2019. 27(3): p. 940–954.e6.

41. Old, W.M., et al., Comparison of Label-free Methods for Quantifying Human Proteins by Shotgun Proteomics*S. Molecular & Cellular Proteomics, 2005. 4(10): p. 1487–1502.

42. Théry, C., et al., Isolation and characterization of exosomes from cell culture supernatants and biological fluids. Curr Protoc Cell Biol, 2006. Chapter 3: p. Unit 3.22.

43. Urbańska, K. and A. Orzechowski, Unappreciated Role of LDHA and LDHB to Control Apoptosis and Autophagy in Tumor Cells. Int J Mol Sci, 2019. 20(9).

44. Zhao, J. and C.J. Zhong, A review on research progress of transketolase. Neurosci Bull, 2009. 25(2): p. 94–9.

45. Novak, A., J.W. Callahan, and J.A. Lowden, Classification of disorders of GM2 ganglioside hydrolysis using 3H-GM2 as substrate. Biochim Biophys Acta, 1994. 1199(2): p. 215–23.

46. Kugeratski, F.G., et al., Quantitative proteomics identifies the core proteome of exosomes with syntenin-1 as the highest abundant protein and a putative universal biomarker. Nature Cell Biology, 2021. 23(6): p. 631–641.

47. Hyenne, V., et al., RAL-1 controls multivesicular body biogenesis and exosome secretion. Journal of Cell Biology, 2015. 211(1): p. 27–37.

48. Colombo, M., et al., Analysis of ESCRT functions in exosome biogenesis, composition and secretion highlights the heterogeneity of extracellular vesicles. Journal of cell science, 2013. 126(24): p. 5553-5565.

49. Zhang, H., et al., Identification of distinct nanoparticles and subsets of extracellular vesicles by asymmetric flow field-flow fractionation. Nature cell biology, 2018. 20(3): p. 332–343.

50. Mizenko, R.R., et al., Tetraspanins are unevenly distributed across single extracellular vesicles and bias sensitivity to multiplexed cancer biomarkers. Journal of Nanobiotechnology, 2021. 19(1): p. 250.

51. Mathieu, M., et al., Specificities of exosome versus small ectosome secretion revealed by live intracellular tracking of CD63 and CD9. Nature Communications, 2021. 12(1): p. 4389.

52. Albacete-Albacete, L., et al., ECM deposition is driven by caveolin-1–dependent regulation of exosomal biogenesis and cargo sorting. Journal of Cell Biology, 2020. 219(11).

53. Provenzano, P.P., et al., Collagen reorganization at the tumor-stromal interface facilitates local invasion. BMC Medicine, 2006. 4(1): p. 38.

54. Dienel, G.A., Chapter 3 - Energy Metabolism in the Brain, in From Molecules to Networks (Third Edition), J.H. Byrne, R. Heidelberger, and M.N. Waxham, Editors. 2014, Academic Press: Boston. p. 53–117.

55. Alfarouk, K.O., et al., The Pentose Phosphate Pathway Dynamics in Cancer and Its Dependency on Intracellular pH. Metabolites, 2020. 10(7).

56. Feng, Y., et al., Lactate dehydrogenase A: A key player in carcinogenesis and potential target in cancer therapy. Cancer Med, 2018. 7(12): p. 6124–6136.

57. Costa-Silva, B., et al., Pancreatic cancer exosomes initiate pre-metastatic niche formation in the liver. Nature cell biology, 2015. 17(6): p. 816–826.

58. Fedele, C., et al., The αvβ6 integrin is transferred intercellularly via exosomes. Journal of Biological Chemistry, 2015. 290(8): p. 4545–4551.

59. Verel-Yilmaz, Y., et al., Extracellular Vesicle-Based Detection of Pancreatic Cancer. Front Cell Dev Biol, 2021. 9: p. 697939.

60. Zhu, K., et al., Single-cell analysis reveals the pan-cancer invasiveness-associated transition of adipose-derived stromal cells into COL11A1-expressing cancer-associated fibroblasts. PLoS Comput Biol, 2021. 17(7): p. e1009228.

61. Ueno, H., et al., Prognostic value of desmoplastic reaction characterisation in stage II colon cancer: prospective validation in a Phase 3 study (SACURA Trial). British Journal of Cancer, 2021. 124(6): p. 1088–1097.

62. Shani, O., et al., Evolution of fibroblasts in the lung metastatic microenvironment is driven by stage-specific transcriptional plasticity. eLife, 2021. 10: p. e60745.

63. Savino, A., et al., Meta-Analysis of Microdissected Breast Tumors Reveals Genes Regulated in the Stroma but Hidden in Bulk Analysis. 2021.

64. Pradhan, R.N., et al., A bird’s eye view of fibroblast heterogeneity: A pan-disease, pan-cancer perspective. Immunol Rev, 2021. n/a(n/a).

65. Hutton, C., et al., Single-cell analysis defines a pancreatic fibroblast lineage that supports anti-tumor immunity. Cancer Cell, 2021.

66. Chen, K., et al., Single-cell RNA-seq reveals dynamic change in tumor microenvironment during pancreatic ductal adenocarcinoma malignant progression. EBioMedicine, 2021. 66: p. 103315.

67. Buechler, M.B., et al., Cross-tissue organization of the fibroblast lineage. Nature, 2021. 593(7860): p. 575–579.

68. Garcia, P.E., et al., Pancreatic Fibroblast Heterogeneity: From Development to Cancer. cells, 2020. 9(11): p. 2464.

69. Schindelin, J., et al., Fiji: an open-source platform for biological-image analysis. Nature Methods, 2012. 9(7): p. 676–682.

70. Clark, K., et al., A specific {alpha}5{beta}1-integrin conformation promotes directional integrin translocation and fibronectin matrix formation. J Cell Sci, 2005. 118(2): p. 291–300.

71. Kielkopf, C.L., W. Bauer, and I.L. Urbatsch, Bradford Assay for Determining Protein Concentration. Cold Spring Harb Protoc, 2020. 2020(4): p. 102269.

72. Cox, J. and M. Mann, MaxQuant enables high peptide identification rates, individualized p.p.b.-range mass accuracies and proteome-wide protein quantification. Nature Biotechnology, 2008. 26(12): p. 1367–1372.

73. Cox, J., et al., Andromeda: A Peptide Search Engine Integrated into the MaxQuant Environment. Journal of Proteome Research, 2011. 10(4): p. 1794–1805.

74. Tyanova, S., et al., The Perseus computational platform for comprehensive analysis of (prote)omics data. Nature Methods, 2016. 13(9): p. 731–740.

