## Supplemental Figures for "NetrinG1^+^ cancer-associated fibroblasts generate unique extracellular vesicles that support the survival of pancreatic cancer cells under nutritional stress"

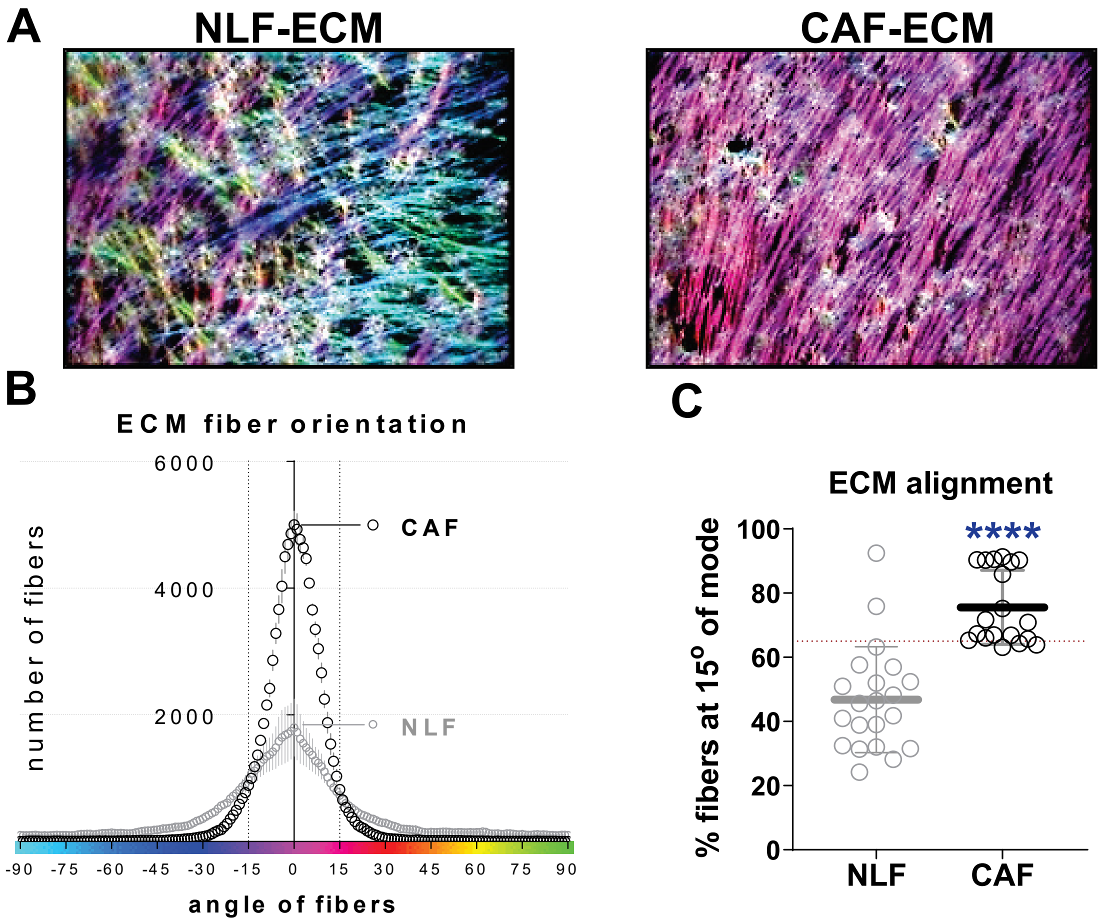


**Supplemental 1: Confirmation that CAFs produce an ECM distinct from NLF in cells.** (A) Fibronectin fiber orientation measured via the orientation-J plugin in FIJI assigns a color gradient based on degree of orientation around a mode angle. (B) Histogram distribution of quantified ECM fiber orientation visualizing fiber alignment. The area denoted between the two dotted lines represents the amount of fibers oriented at 15° angles from the mode (normalized to 0°). (C) Percentage of fibers within 15° of the mode (obtained from the dotted line area of panel B) were compared to question significance in alignment. Red dotted line denotes 65% and represents threshold for determining bona-fide CAF activation (e.g., alignment). n=3. Bars = standard error. Statistical test used: Mann-Whitney t-test.

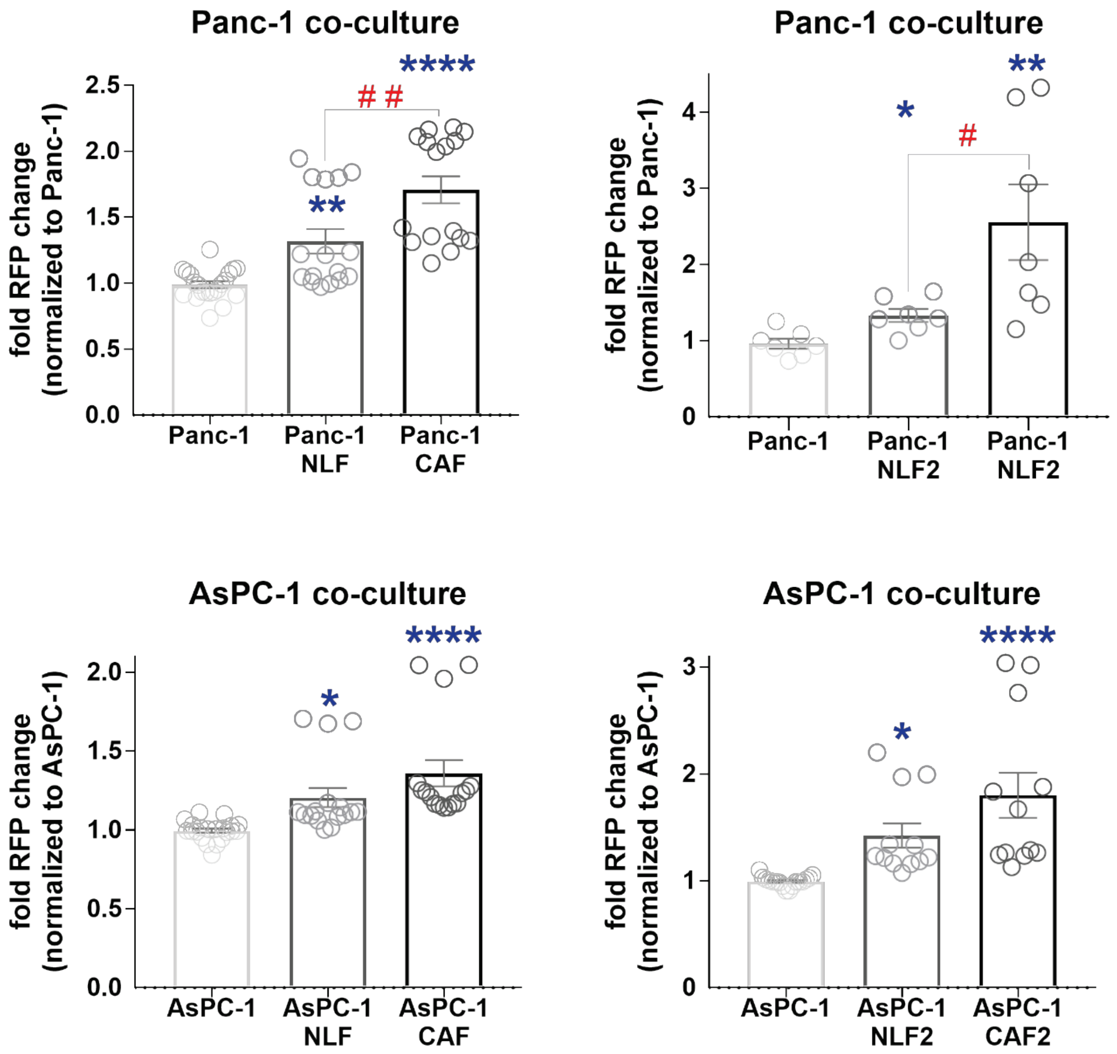


**Supplemental 2: Direct co-culture of CAFs support PDAC cells during nutrient-deprivation.** Direct co-culture of PDAC cells (Panc-1 A-B; AsPC-1 C-D) and CAF and NLF fibroblastic cells harvested from two independent patients. n=3. Bars = standard error. * denotes comparisons between experimental and PBS-treated condition (negative control), while # denotes comparisons between experimental conditions linked by connecting lines. Statistical test used: 1-way ANOVA, with multiple comparisons using the Tukey correction

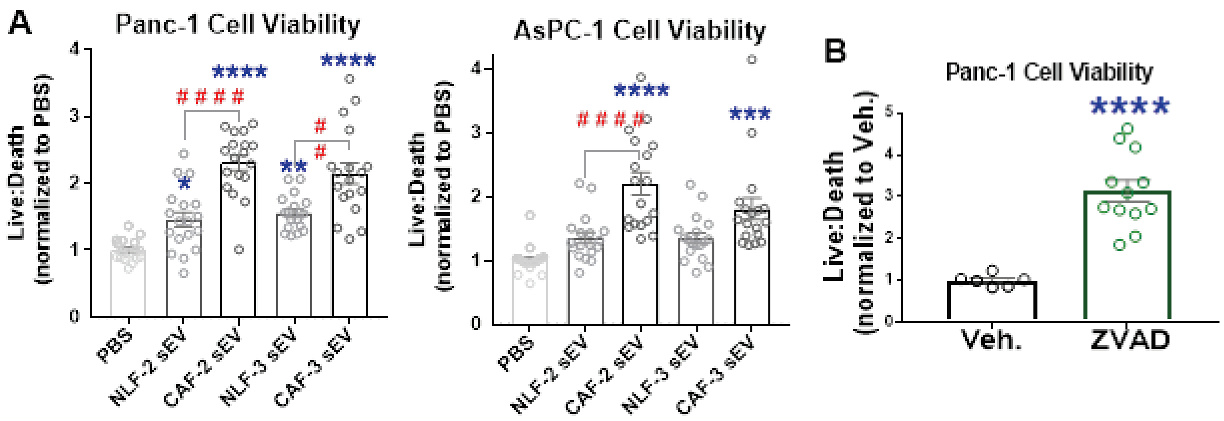


**Supplemental 3: Paracrine CAF-mediated support of nutrient-deprived PDAC.** (A) Viability assay of PANC-1 (left) and AsPC-1 (right) showing that two added CAFs and NFLs (e.g., obtained from different patients) present with similar survival benefits (or lack thereof) as the fibroblastic cells used in main figure 3. n=3 biological replicates; each biological replicate contains 6 technical repeats. Technical repeats for each biological replicate were normalized to the PBS-treated condition. Bars = standard error. Statistical test used: 1-way ANOVA, with multiple comparisons using the Tukey correction. * denotes comparison to PBS (negative control), while # denotes comparison between conditions linked by connecting lines. (B) PANC-1 cell viability assay designed to assess whether cells die due to apoptosis via treatment with 1 μM Z-VAD-FMK (ZVAD). Note the increased survival of PDAC cells by 3-fold, compared to vehicle treated cells (Veh.) cultured in CM. n=3. Bars = standard error.

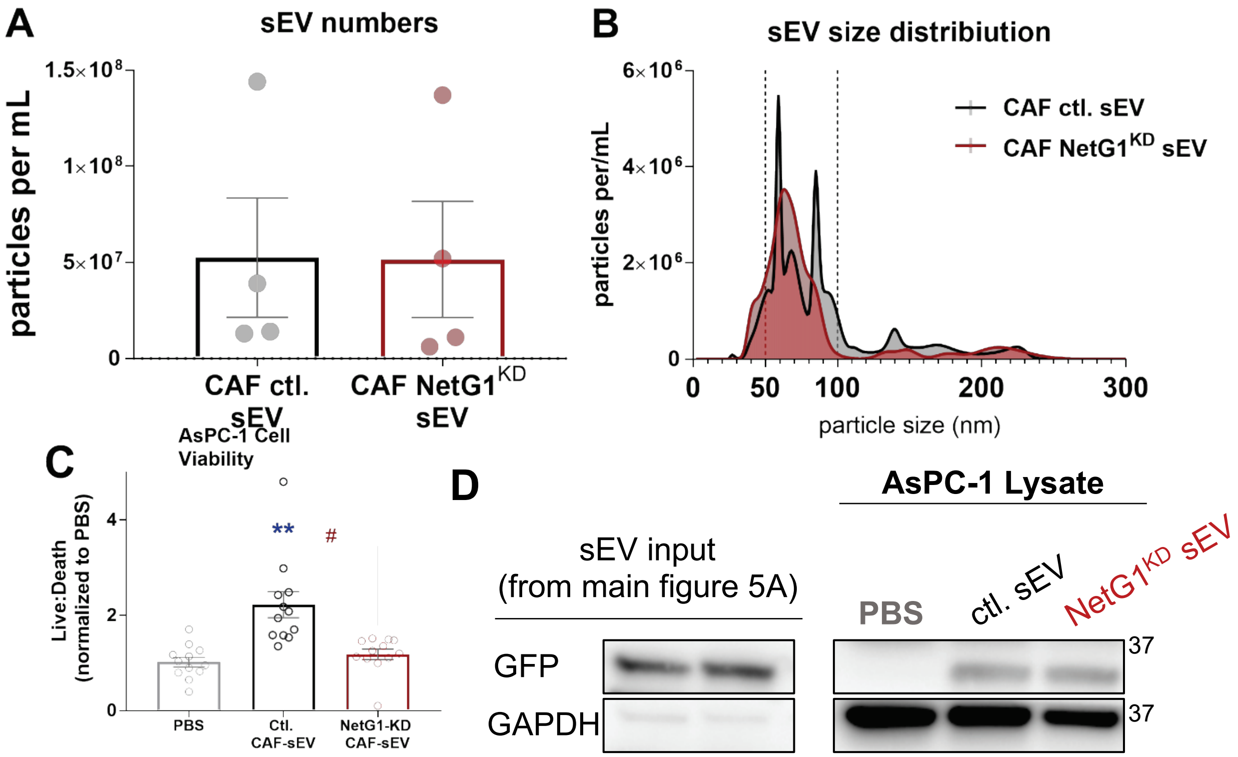


**Supplemental 4: NetG1 manipulation in CAFs alters EV size distribution and pro-tumor function.** (A) Particle concentrations assessed using the nanosight platform. n=4. Bars = standard error. Statistics = Welch's t-test. (B) Histogram of nanoparticle distribution of Ctl.CAF and NetG1-KD CAFs sEV fractions, superimposed, using the NanoSight platform. (C and E) Viability assays of AsPC-1 cells measuring live-to-dead cell ratios. n=4 biological replicates; each biological replicate contains 6 technical repeats. Repeats per replicate were normalized to PBS. Bars = standard error. * denotes comparison to PBS (negative control), while # denotes comparison between experimental conditions linked by connecting lines. Statistics = 1-way ANOVA, with multiple comparisons using the Tukey correction. (D) Representative Western Blots showing input values of GFP in sEVs and amount of GFP incorporated by PDAC (AsPC-1) cell lysates following 24 hours post treatment.

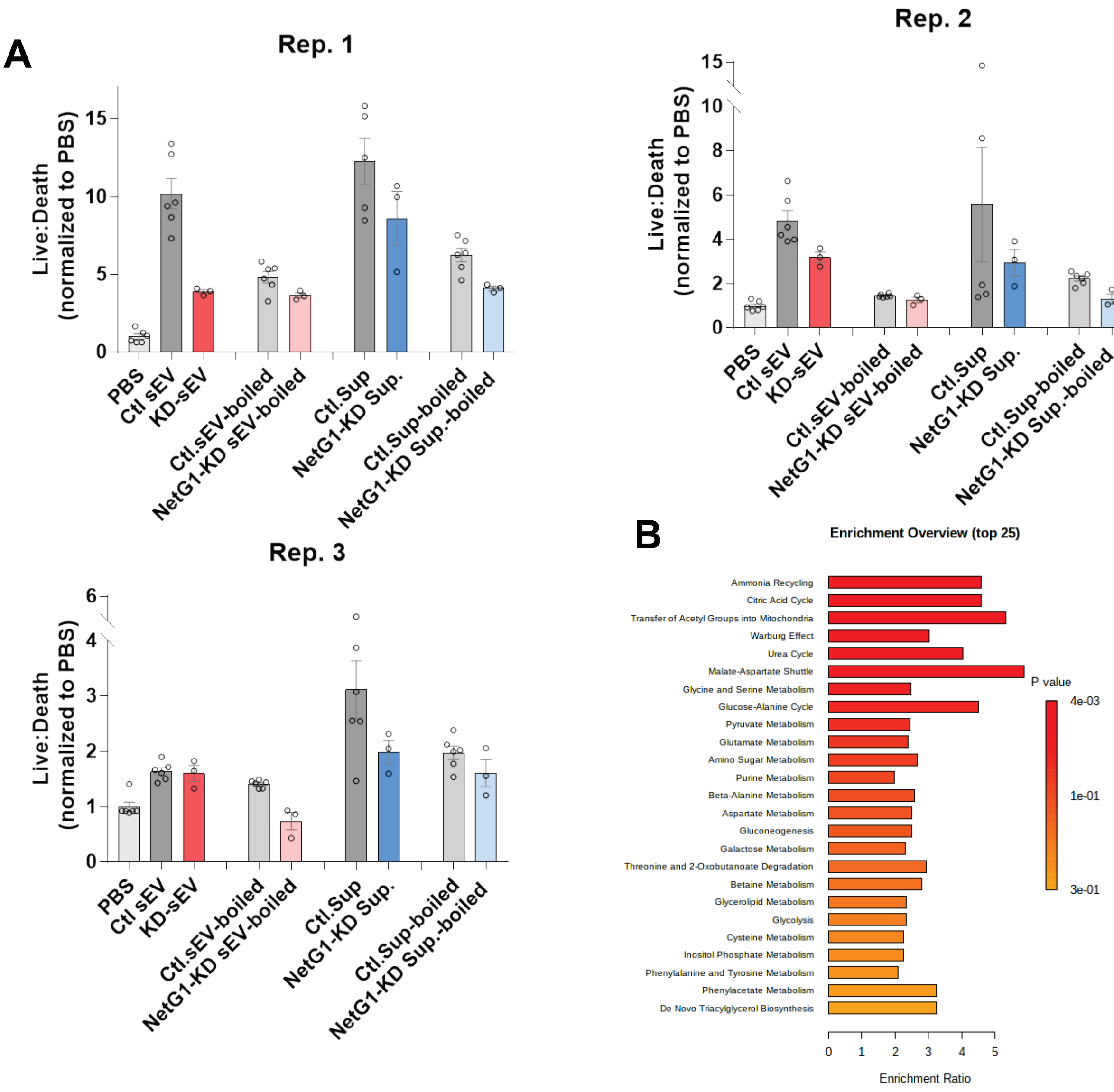


(A) sEV and supernatant fractions collected from control CAFs (ctl.) or NetG1-KD CAFs (NetG1-KD) were boiled and filtered prior to testing their ability to sustain PDAC cells survival under nutritional stress. Graphs represent three repetitions (Rep) (e.g. independent experiments). (B) metabolomics data was queried using the MetaboAnalyst and the most significant pathways enriched in sEVs from control CAFs compared to sEVs collected from NetG1-KD CAFs are shown.

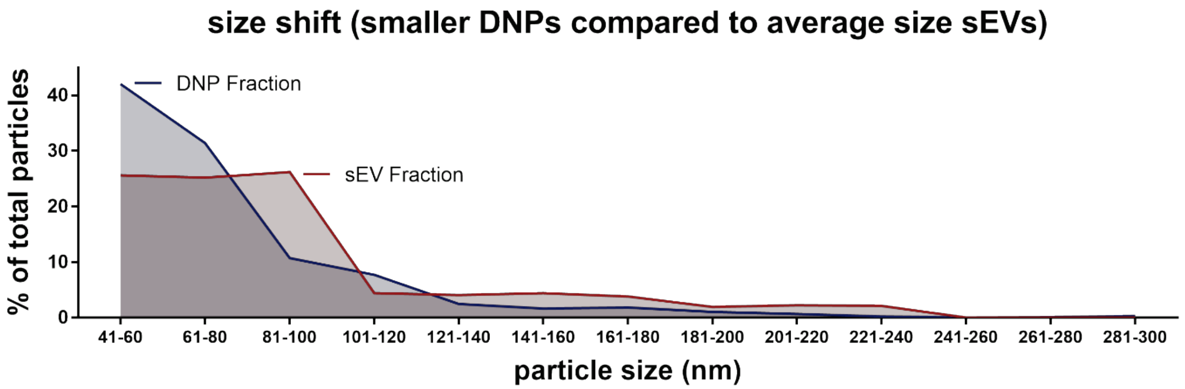


**Supplemental 6: DNP pellet is enriched with sub-exosomal sized EVs**.
Representative nanoparticle distribution of CAF sEV (red) and DNP (blue) fractions. Graphs represent size distributions (size range divided by the total amount of particles per sample read to denote percentages) using the NanoSight platform. Size had a cut off value of 40 nm.

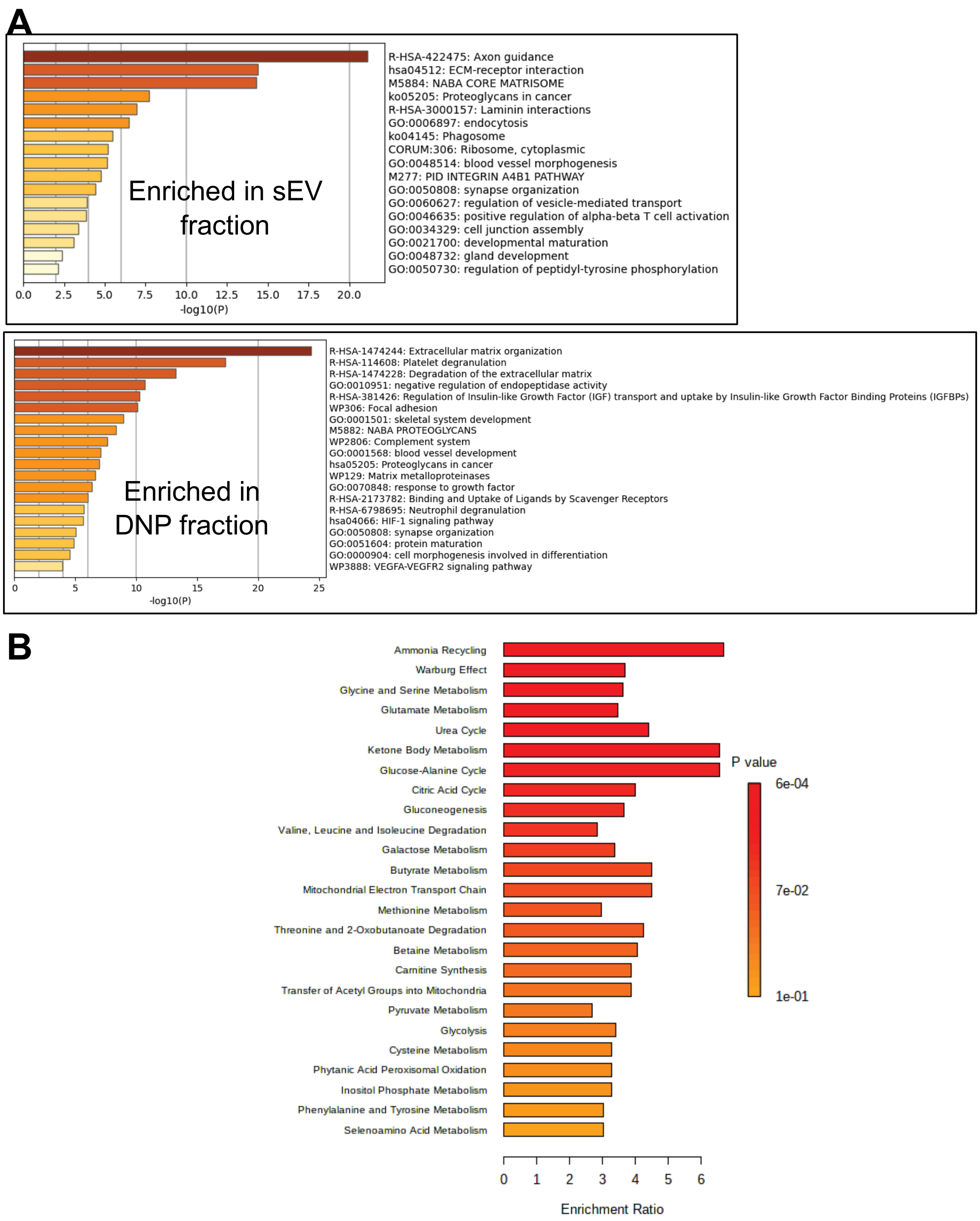


**Supplemental 7: Enriched gene ontology clusters from proteomic and metabolomic analysis comparing sEV and DNP fractions.**

(A) Following LFQ experimental analysis, all proteins whose p-values indicated significantly enriched levels in sEVs (top) or DNP (bottom) were identified for enriched terms including GO/KEGG terms, canonical pathways, and hallmark gene sets. Ontology clusters were sorted, calculated, and ranked by p-values using the Metascape.org gene annotation and analysis resource. (B) Metabolomics data comparing sEVs to DNPs was queried using the MetaboAnalyst and the top 25 significant pathways enriched in sEVs are shown.
