## Supplementary material for "NetrinG1^+^ cancer-associated fibroblasts generate unique extracellular vesicles that support the survival of pancreatic cancer cells under nutritional stress": Key resources table

Reagent Table

| Reagent | Source | Identifier |
|---|---|---|
| **Antibodies** | | |
| Alix (1A12) | Santa Cruz | Cat# sc-53540, RRID:AB_673819 |
| CD63 (MX-49.129.5) | SantaCruz | Cat# sc-5275, RRID:AB_627877 |
| CD81 (B-11) | Santa Cruz | Cat# sc-166029, RRID:AB_2275892 |
| Fibronectin antibody | Sigma-Aldrich | Cat# F0791, RRID:AB_476961 |
| GAPDH | Abcam | Cat# ab9485, RRID:AB_307275 |
| GFP (D5.1) XP | Cell Signaling | Cat# 2956, RRID:AB_1196615 |
| Hexokinase 1 (G-1) | Santa Cruz | sc-46695, RRID:AB_627721) |
| Histone 3 | Cell Signaling | Cat# 9715, RRID:AB_331563 |
| Integrin alpha 5 | Santa Cruz | sc-10729 |
| Netrin G1 Ligand [N1N3] | Genetex | Cat# GTX121508, RRID:AB_10721992 |
| NetrinG1 | Santa Cruz | Cat# sc-271774, RRID:AB_10707668 |
| PARP | Cell Signaling | Cat# 9542, RRID:AB_2160739 |
| SNAKA51 Antibody | Gift from M Humphries | N/A |
| TSG101 [EPR7130(B] | Abcam | Cat# ab125011, RRID:AB_10974262 |
| αSMA antibody | Sigma-Aldrich | Cat# A2547, RRID:AB_47670 |
| Anti-Mouse IgG HRP | Cell signaling | Cat# 7076, RRID:AB_330924 |
| Anti-Rabbit IgG HRP | Cell signaling | Cat# 7074, RRID:AB_2099233 |
| Cy™5 AffiniPure Donkey Anti-Rabbit IgG (H+L) | Jackson ImmunoResearch | Cat# 711-175-152, RRID:AB_2340607 |
| Rhodamine Red™-X (RRX) AffiniPure Donkey Anti-Mouse IgG (H+L) | Jackson ImmunoResearch | Cat# 715-295-151, RRID:AB_2340832 |
| SYBR Green I | Thermo Scientific | Cat# S7563 |
| **Chemicals** | | |
| Protein Standard | BioRad | 1610374 |
| 4x Laemmli Sample Buffer | BioRad | 161-0747 |
| Quick Start Bradford 1x Dye Reagent | BioRad | 500-0205 |
| Odyssey Blocking Buffer (PBS) | Li-COR Biosciences | 927-70001 |
| 2-mercaptoethanol | Sigma-Aldrich | M3148-100ML |
| L-Ascorbic acid | Sigma-Aldrich | A92902-100G |
| Tween 20 | Fischer Science | BP337-500 |
| 25% Glutaraldehyde Solution in water | Sigma-Aldrich | G6257-1L |
| Paraformaldehyde 16% solution, EM grade | Electron Microscopy Sciences | 15710 |
| Uranyl Acetate 2% solution | Electron Microscopy Sciences | 22400-2 |
| Ethanolamine | Sigma-Aldrich | E9508-1L |
| Dimethyl sulfoxide-for HPLC | Sigma-Aldrich | 34869 |
| ProLong™ Gold Antifade | Invitrogen | P36930 |

| | | |
|---|---|---|
| Mountant | | |
| Sytox Blue | Invitrogen | S11348 |
| Protease inhibitor | Thermo Scientific | 36978 |
| Phosphatase inhibitor | Thermo Scientific | A32957 |
| **Cell Culture** | | |
| FBS | Peak Serum | PS-FB2 |
| DMEM (High Glucose + Phenol Red) | Cell Culture Facility (FCCC) | N/A |
| DMEM (low Glucose+ Phenol Red) | Cell Culture Facility (FCCC) | N/A |
| DMEM (High glucose – Phenol Red) | Corning | 17-205-CV |
| DMEM (low glucose – Phenol red) | Thermo Scientific | 11054001 |
| M3 Media Supplement | Incell | M300A |
| L-Glutamine | Corning | #25-005CI |
| Penicillin/Streptomycin | Corning | #30-002-CI |
| **Cell Lines** | | |
| Panc-1 | ATCC | RRID: CVCL_0480 |
| AsPC-1 | ATCC | CRL-1682 |
| Cancer associated fibroblasts (patient derived) | Fox Chase Cancer Center | N/A |
| Tumor adjacent fibroblasts (patient derived) | Fox Chase Cancer Center | N/A |
| **Materials and Machines** | | |
| Autoradiography Film | Midwest Scientific | BX810 |
| Film Developer | AFP imaging | Mini-Medical Series |
| Auto Fluorescent imager FluroChem E | Protein Simple | 3184363 |
| Confocal Microscope | Nikon | Eclipse Ti2 |
| Microscope Camera Orca-flash 4.0 | Hamamatsu | C11440 |
| 0.22 micron PVDF syringe filter (33 mm) | Millipore | SLGV033RS |
| Polypropylene Ultracentrifuge tubes (11 x 34 mm) | Beckman Coulter | 347357 |
| Polypropylene Quick-Seal tubes (11 x 32 mm) | Beckman Coulter | 344625 |
| Amicon Ultra Centrifuge Filter Tubes | Millipore | UFC500324 |
| Electron Microscopy Formvar coated Nickel (300 mesh) grid | Electron Microscopy Sciences | 24928 |
